# Genome-wide single-molecule analysis of long-read DNA methylation reveals heterogeneous patterns at heterochromatin

**DOI:** 10.1101/2022.11.15.516549

**Authors:** Lyndsay Kerr, Ioannis Kafetzopoulos, Ramon Grima, Duncan Sproul

## Abstract

High-throughput sequencing technology is central to our current understanding of the human methylome. The vast majority of studies use chemical conversion to analyse bulk-level patterns of DNA methylation across the genome from a population of cells. While this technology has been used to probe single-molecule methylation patterns, such analyses are limited to short reads of a few hundred basepairs. DNA methylation can also be directly detected using Nanopore sequencing which can generate reads measuring megabases in length. However, thus far these analyses have largely focused on bulk-level assessment of DNA methylation. Here, we analyse DNA methylation in single Nanopore reads with a mean length of 24.6kb, to show that bulk-level metrics underestimate large-scale heterogeneity in the methylome. We use the correlation in methylation state between neighbouring sites to quantify single-molecule heterogeneity and find that heterogeneity varies significantly across the human genome, with some regions having heterogeneous methylation patterns at the single-molecule level and others possessing more homogeneous methylation patterns. By comparing the genomic distribution of the correlation to epigenomic annotations, we find that the greatest heterogeneity in single-molecule patterns is observed within heterochromatic partially methylated domains (PMDs). In contrast, reads originating from euchromatic regions and gene bodies have more ordered DNA methylation patterns. By analysing the patterns of single molecules in more detail, we show the existence of a 185bp periodicity in DNA methylation that accounts for some of the heterogeneity we uncover in long single-molecule DNA methylation patterns. We find that this periodic structure is partially masked in bulk data in a manner that is consistent with imperfect phasing of nucleosomes between molecules. Our findings demonstrate the power of single-molecule analysis of long-read data to understand the structure of the human methylome.

## Introduction

DNA methylation is an epigenetic mark commonly associated with gene repression [1]. In mammals it is prevalent on cytosines within CpG dinucleotides, with 70 − 80% of CpGs being methylated in most mammalian cells [2].

High-throughput sequencing approaches have resulted in the description of many human methylomes using bisulfite conversion in combination with whole-genome sequencing (Whole-Genome Bisulfite Sequencing, WGBS) [3, 4]. Analysis of WGBS data has confirmed the pervasive nature of CpG methylation in human genomes and uncovered differences in the DNA methylation patterns between different cell types and tissues [5, 6]. They have also detailed differences in DNA methylation occurring in human disease, particularly cancer [7].

These insights have arisen from the analysis of bulk DNA methylation patterns. High-throughput bisulfite sequencing data can also be analysed at the single-read level to understand single-molecule heterogeneity in DNA methylation patterns. Analyses of single-read patterns have been used to quantify the degree of inter-molecular methylation heterogeneity in human samples [8, 9]. Further analyses have quantified differences in this heterogeneity between normal and cancer cells using information theory [10]. DNA methylation heterogeneity has been shown to change over time in cell culture [11] and in acute myeloid leukemia upon relapse [12]. Inter-molecular methylation heterogeneity has been attributed to epi-polymorphisms [8] and allele-specific methylation [13]. The mixture of cell types in a sample is also an important source of DNA methylation heterogeneity [14, 15], that could explain part of the inter-molecular heterogeneity seen in these analyses.

However, the degradation of DNA caused by bisulfite conversion [16] and the widespread use of Illumina sequencing technology limits single-molecule analyses of WGBS data to short regions of a few hundred basepairs. The limited number of CpGs captured in these reads restricts analyses of intra-molecular DNA methylation heterogeneity. Nanopore sequencing enables the generation of reads over a megabase in length [17], four orders of magnitude longer than those assayed by WGBS. DNA modifications including 5-methylcytosine can also be directly detected in these reads without chemical conversion [18]. This potentially enables the analysis of molecular heterogeneity in DNA methylation patterns at a far greater scale than previously examined.

Here we conduct a genome-wide analysis of single-molecule DNA methylation patterns in long reads derived from Nanopore sequencing in order to understand the nature of large-scale intra-molecular DNA methylation heterogeneity in the human genome. Our work demonstrates that intra-molecular DNA methylation heterogeneity is abundant in heterochromatin, which possesses oscillatory methylation patterns that are partially masked in bulk data.

## Results

### Large-scale methylation heterogeneity is underestimated by bulk analysis of methylomes

To understand the nature of large-scale heterogeneity in DNA methylation patterns, we analysed Nanopore sequencing data derived from GM24385 lymphoblastoid cells. This consisted of 6, 289, 480 reads aligned to autosomes with a mean length of 24.6kb.

We first asked how large-scale intra-molecular DNA methylation heterogeneity varied across single reads. We therefore calculated the mean level of methylation for all aligned single reads along with the coefficient of variation and correlation to capture intra-molecular variation (Fig 1**a**). To focus our attention on large-scale DNA methylation patterns, and reduce noise associated with calculating statistics from a low number of CpGs, only aligned reads with methylation called for ≥ 100 CpG sites were considered (2, 908, 181). All of these single-read measurements varied considerably across aligned reads suggesting that substantial large-scale intra-molecular DNA methylation heterogeneity exists in human cells (Fig 1**b-d**).

**Fig 1.**
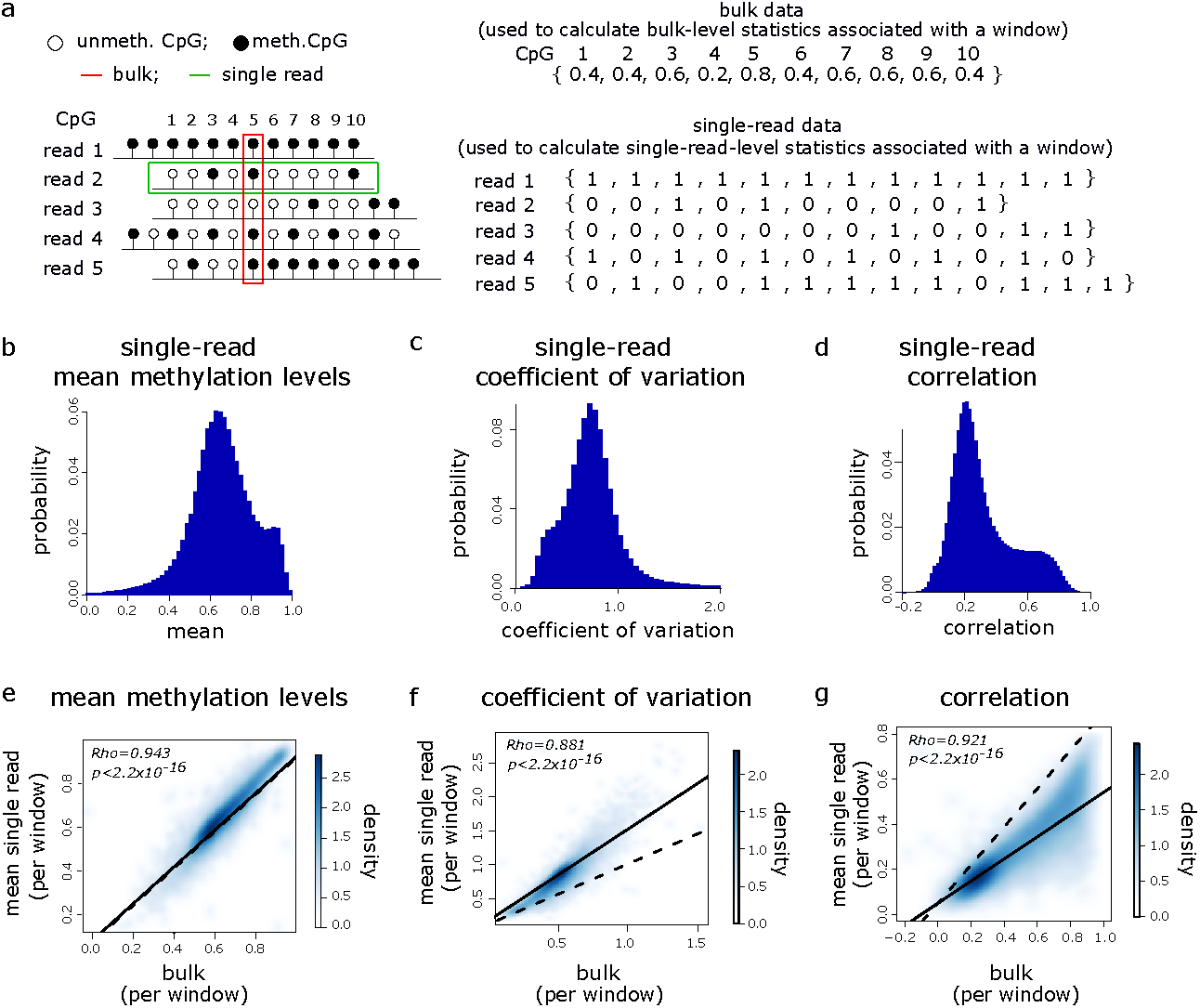
Large-scale methylation heterogeneity is underestimated by bulk analysis of methylomes. **a** Left: schematic of analysis approach. Example single-read (green box) and bulk (red box) analysis for 5 hypothetical reads overlapping 10 common CpGs are shown. Right: the bulk methylation level of a CpG is its number of methylated observations divided by its total number of unmethylated and methylated observations. The red box highlights information used to calculate the bulk methylation level of CpG 5. Single-read data is obtained by creating a vector, *v*, for each read, where *v*_*i*_ = 0, 1 if CpG *i* in the read is unmethylated or methylated respectively. The green box highlights information used to construct *v* for read 2. **b–d** Histograms showing the distributions of single-read statistics for the 2, 908, 181 reads with methylation states for ≥100 CpGs. (**b**) mean methylation levels, (**c**) coefficients of variation and (**d**) correlations between neighbouring CpGs. **e–g** Density scatter plots of mean single-read vs. bulk statistics in 100kb genomic windows (*n* = 26, 687). (**e**) mean methylation levels, (**f**) coefficients of variation and (**g**) correlations between neighbouring CpGs. Dashed and solid lines show lines of identity and linear models fitted to the data respectively. Spearman correlations are shown along with p-values from paired t-tests comparing bulk to mean single-read statistics.

We then asked how these single-read measurements compared to bulk-level measurements (Fig 1**a**). To facilitate this comparison, we segmented the autosomal genome into 100kb non-overlapping windows and calculated the mean of each of the single-read statistics for each window using the reads that aligned entirely within that windows (total of 1, 707, 118 reads for all windows). We compared these mean single-molecule measures to bulk-level statistics for the same windows. We calculated the bulk statistical measure for a window by first calculating the mean methylation level of each CpG in the window using all reads in which it had a defined methylation state. We then calculated the bulk statistics using these mean CpG methylation values. We hypothesised that the bulk and single-read approaches would provide similar estimates of mean methylation levels but differ in their quantification of variation. Measurements of the mean single-read methylation levels in the genomic windows were significantly correlated with bulk mean levels of DNA methylation within the same windows (Fig 1**e**, Spearman correlation = 0.943, *p <* 2.20 × 10^*−*16^). While a two-sided paired t-test indicated significant difference in the bulk means compared to the single-read means, the absolute relative difference of the single-read means compared to the bulk means was small (mean absolute relative difference of 0.0375). As the single-read and bulk estimates of mean methylation are mathematically equivalent, these small differences are likely to be caused by slightly different read and CpG exclusion criteria between the two approaches. As expected, these results suggest that single-read measurements capture largely the same information regarding overall methylation levels as bulk measurements.

We next considered how single-read measurements of heterogeneity compared to their bulk counterparts. Like mean methylation levels, the mean single-read and bulk measurements of the coefficient of variation and correlation were significantly correlated (Fig 1**f, g**, Spearman correlation = 0.881 and 0.921, respectively, *p <* 2.20 × 10^*−*16^ in both cases). However, mean single-read coefficients of variation observed in the genomic windows were significantly greater than that of the equivalent bulk measurements (Fig 1**f**, paired t-test p *<* 2.20 × 10^*−*16^, mean absolute relative difference of 0.338) and a linear model fitted to the two deviated from the line of identity (Fig 1**f** solid vs. dashed line). We also observed that mean single-read correlations were significantly lower than that of the bulk measurements (Fig 1**g**, paired t-test p *<* 2.20 × 10^*−*16^, mean absolute relative difference of 0.621) and a linear model fitted to the two deviated from the line of identity (Fig 1**g** solid vs. dashed line). To ensure that these results were not specific to the window size used, we repeated the analysis using 50kb and 200kb genomic windows observing similar results as when 100kb windows were used (Fig. S1).

These comparisons of single-molecule to bulk measurements suggest that the analysis of DNA methylation in single Nanopore reads captures large-scale intra-molecular DNA methylation heterogeneity that is missed by analysis of bulk DNA methylation patterns.

### Intra-molecular heterogeneity varies across the human genome

Having established that significant large-scale intra-molecular DNA methylation heterogeneity exists in the human methylome, we next asked how this heterogeneity was organised across the genome.

Many measures previously used to quantify single-molecule methylation heterogeneity from short-read data [19] are difficult to apply to long reads because the number of possible states increases with the number of CpGs in the read. We therefore measured single-read methylation heterogeneity using the correlation in methylation state between nearest-neighbour CpGs (see Eq (4)). We chose to measure single-molecule heterogeneity using the correlation rather than the coefficient of variation since the latter does not take into account information relating to the methylation states of neighbouring CpGs. A high correlation indicates high agreement in the methylation state of nearest-neighbour CpGs and thus low intra-molecular heterogeneity. On the other hand, a low correlation indicates high disagreement in the methylation state of nearest-neighbour CpGs and thus high intra-molecular heterogeneity.

To ask how intra-molecular heterogeneity varied across the genome, we calculated the correlation associated with each of the 2, 908, 181 reads containing methylation information for ≥ 100 CpGs that mapped to autosomes. The resulting distribution of individual read correlations was weakly bimodal (Fig 2**a**). We observed a significantly different unimodal distribution (Kolmogorov-Smirnov test, *p <* 2.20 × 10^*−*16^) characterised by lower correlation values when we calculated the correlation using reads simulated to have a random intra-molecular methylation pattern (Fig 2**a**, see methods). These results suggest that reads had methylation patterns that were non-random and that intra-molecular methylation heterogeneity might vary by location in the genome.

**Fig 2.**
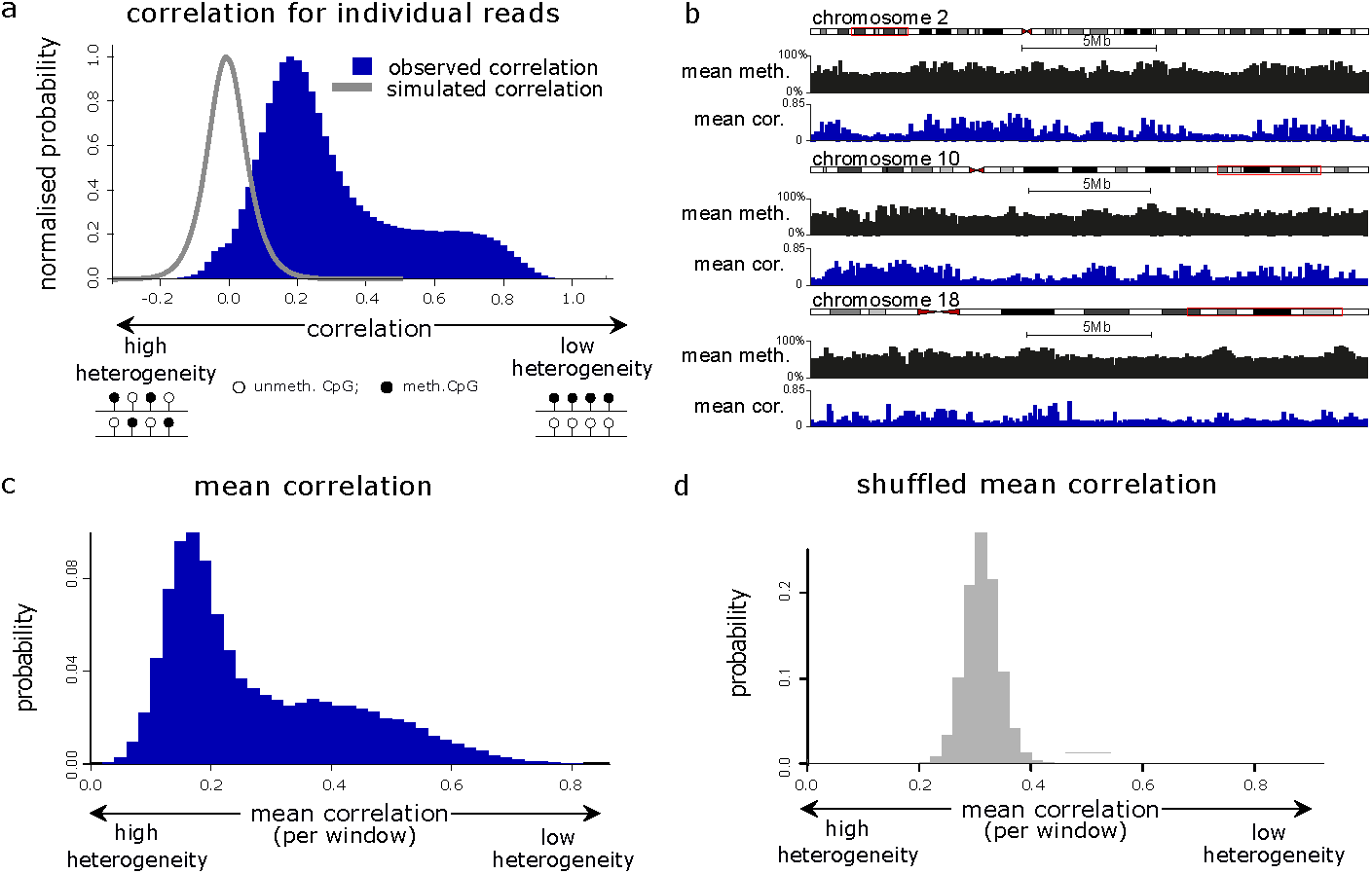
Intra-molecular heterogeneity varies across the human genome. **a** Histogram showing the correlation distribution for the 2, 908, 181 reads containing methylation states for ≥ 100 CpGs. This is compared to the correlation calculated from reads simulated to have a random intra-molecular methylation pattern (grey line). **b** Genome browser plots showing the mean methylation levels and mean correlation in 100kb windows for selected genomic windows. **c** Histogram of the mean correlation distribution for 100kb genomic windows (*n* = 26, 951), calculated using reads that align entirely within each window. **d** Histogram showing the mean shuffled correlation in 100kb windows (*n* = 26, 951), calculated using random reads rather than those that align to each window.

To understand whether the degree of methylation heterogeneity varied by location in the genome, we examined the mean correlation across reads entirely aligned within 100kb windows (Fig 2**b**, [20]. We then compared the distribution of mean correlation to that obtained when we assigned random reads to each genomic window (shuffled correlation). The distribution of mean correlation for genomic windows was bimodal and significantly different from the unimodal distribution observed for the shuffled correlation (Kolmogorov-Smirnov test, *p <* 2.20 × 10^*−*16^, Fig 2**c, d**). Moreover the peak of the unimodal shuffled correlation distribution lies between the two peaks associated with the bimodal mean correlation distribution. This is consistent with some regions of the human genome possessing a lower degree of intra-molecular heterogeneity than would be expected by chance, and other regions displaying a higher degree of intra-molecular heterogeneity.

To validate that intra-molecular methylation heterogeneity varies across the genome, we conducted a similar analysis using an alternative measure of intra-molecular heterogeneity, the Read Transition Score (RTS; [21], Eq. 11, Fig. S2**a**). The RTS measures the probability that two nearest-neighbour CpGs have different methylation states and ranges from 0 to 1. An RTS of 1 indicates that no nearest-neighbour CpGs have the same methylation state and an RTS of 0 indicates that only a single methylation state is observed in the read (Fig. S2**a**). Hence we take a low RTS to be indicative of low heterogeneity within the single-molecule pattern and a high RTS to be indicative of high heterogeneity within the single-molecule pattern. The RTS for individual reads was again observed to be bimodally distributed (Fig. S2**b**) and showed significant negative correlation with the single-read correlation (Spearman correlations = − 0.666, *p <* 2.20 × 10^*−*16^, Fig. S2**c**). Similarly, the mean RTS in 100kb windows showed significant negative correlation with the mean correlation in 100kb windows (Spearman correlations = − 0.768, *p <* 2.20 × 10^*−*16^, Fig. S2**d**). In addition, the distribution of mean RTS observed in 100kb windows was significantly different from that obtained from shuffled reads, consistent with the degree of intra-molecular heterogeneity varying across the human genome (Fig. S2**e**,**f**, Kolmogorov-Smirnov test, *p <* 2.20 × 10^*−*16^).

Taken together, these analyses suggest that the degree of large-scale intra-molecular DNA methylation heterogeneity varies between different regions of the human genome.

### Intra-molecular and inter-molecular methylation heterogeneity co-occur in the human genome

Intra-molecular methylation heterogeneity can occur in a manner that is consistent or inconsistent across reads arising from the same region of the genome. This means that a low correlation in the methylation state between nearest-neighbour CpGs could be compatible with either high or low inter-molecular methylation heterogeneity (Fig. S3).

To understand the degree of inter-molecular heterogeneity associated with regions of the genome displaying high intra-molecular methylation heterogeneity, we computed the measure *d*_*i*_. This quantifies the degree of inter-molecular heterogeneity across reads for a given CpG as *d*_*i*_ = |*m*_*i*_ − 0.5| for a CpG *i* with methylation level *m*_*i*_. This ranges from 0, which indicates maximum inter-read heterogeneity when the unmethylated and methylated states are observed equally across reads, to 0.5, when a CpG has a uniformly methylated or unmethylated state.

We then calculated *d*_*i*_ for every CpG in the genome before comparing the values observed for regions of the genome associated with the highest or lowest degree of intra-read methylation heterogeneity as measured by the correlation. The distribution of *d*_*i*_ values for the 10% of 100kb genomic windows with the highest mean correlation were significantly skewed towards high values of *d*_*i*_ in comparison to the background of all genomic windows (Fig 3**a**,**b**, *p <* 2.20 × 10^*−*16^, Kolmogorov-Smirnov test). This indicates that these regions display low inter-read heterogeneity as well as low intra-read heterogeneity. In contrast the *d*_*i*_ scores of CpGs found in the 10% of 100kb genomic windows with the lowest mean correlation were significantly lower than the background of all genomic windows (Fig 3**a**,**c**, *p <* 2.20 × 10^*−*16^, Kolmogorov-Smirnov test) suggesting that they also show a greater degree of inter-read heterogeneity.

**Fig 3.**
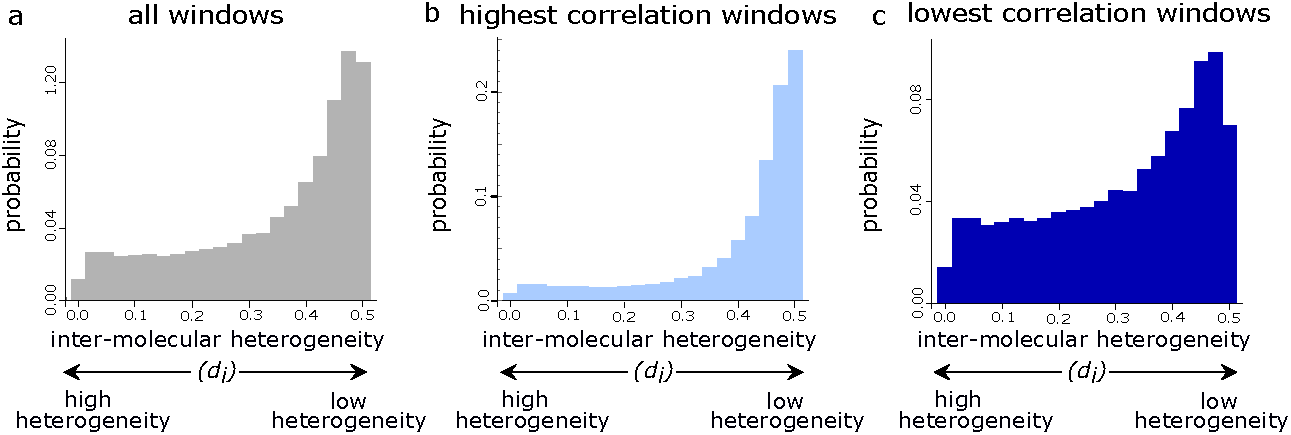
Intra-molecular and inter-molecular methylation heterogeneity co-occur in the human genome. **a–c** Histograms showing the distributions of *d*_*i*_ (**a**) for the 26, 205, 980 CpGs in all windows, (**b**) for the 4, 504, 144 CpGs in the 10% of windows with the highest correlation and (**c**) for the 1, 535, 028 CpGs in the 10% of windows with the lowest correlation.

These analyses of CpG methylation levels demonstrate that regions of the genome with a high degree of intra-molecular DNA methylation heterogeneity also posses high inter-molecular methylation heterogeneity.

### Heterochromatin possesses heterogeneous methylation patterns

To understand the nature of the regions of the human genome which had the greatest degree of intra-molecular DNA methylation heterogeneity, we asked whether they were associated with particular genomic annotations or chromatin states.

We first analysed the 10% of 100kb genomic windows with the lowest mean correlation (i.e. the 10% most heterogeneous windows). We observed that these genomic windows were significantly depleted in CpG islands compared to the background of all genomic windows (fold-enrichment of 0.0154, Wilcoxon test, *p <* 2.20 × 10^*−*16^) but showed no significant depletion or enrichment in genes (fold-enrichment of 0.936, Wilcoxon test, *p* = 0.566). The least heterogeneous genomic windows (the 10% with the highest mean correlation), were significantly enriched in both genes and CpG islands (fold-enrichments of 1.21 and 4.18, Wilcoxon tests *p* = 1.09 × 10^*−*6^ and *p <* 2.20 × 10^*−*16^, respectively).

To further understand the characteristics of regions displaying high methylation heterogeneity, we cross-referenced them to chromatin state data generated by the ENCODE project [22, 23]. These projects have used hidden Markov models to partition the genome into distinct chromatin states (chromHMM). Consistent with their observed enrichment in genes and CpG islands, the least heterogeneous genomic windows were significantly enriched in chromHMM-defined promoters, enhancers and transcribed regions from GM12878 lymphoblastoid cells (Fig 4**a**, Wilcoxon tests with *p <* 2.20 × 10^*−*16^ in each case). They were also significantly enriched in insulators and polycomb-repressed regions (Fig 4**a**, Wilcoxon tests, *p* = 6.03 × 10^*−*12^ and *p <* 2.20 × 10^*−*16^, respectively). In contrast, the most heterogeneous windows were significantly enriched in chromHMM-defined heterochromatic regions (Fig 4**b**, Wilcoxon test with *p <* 2.20 × 10^*−*16^). We observed similar enrichment results when the analysis was repeated using 50kb and 200kb genomic windows (Fig. S4**a, b**) suggesting that our observations were independent of the choice of genomic window size.

**Fig 4.**
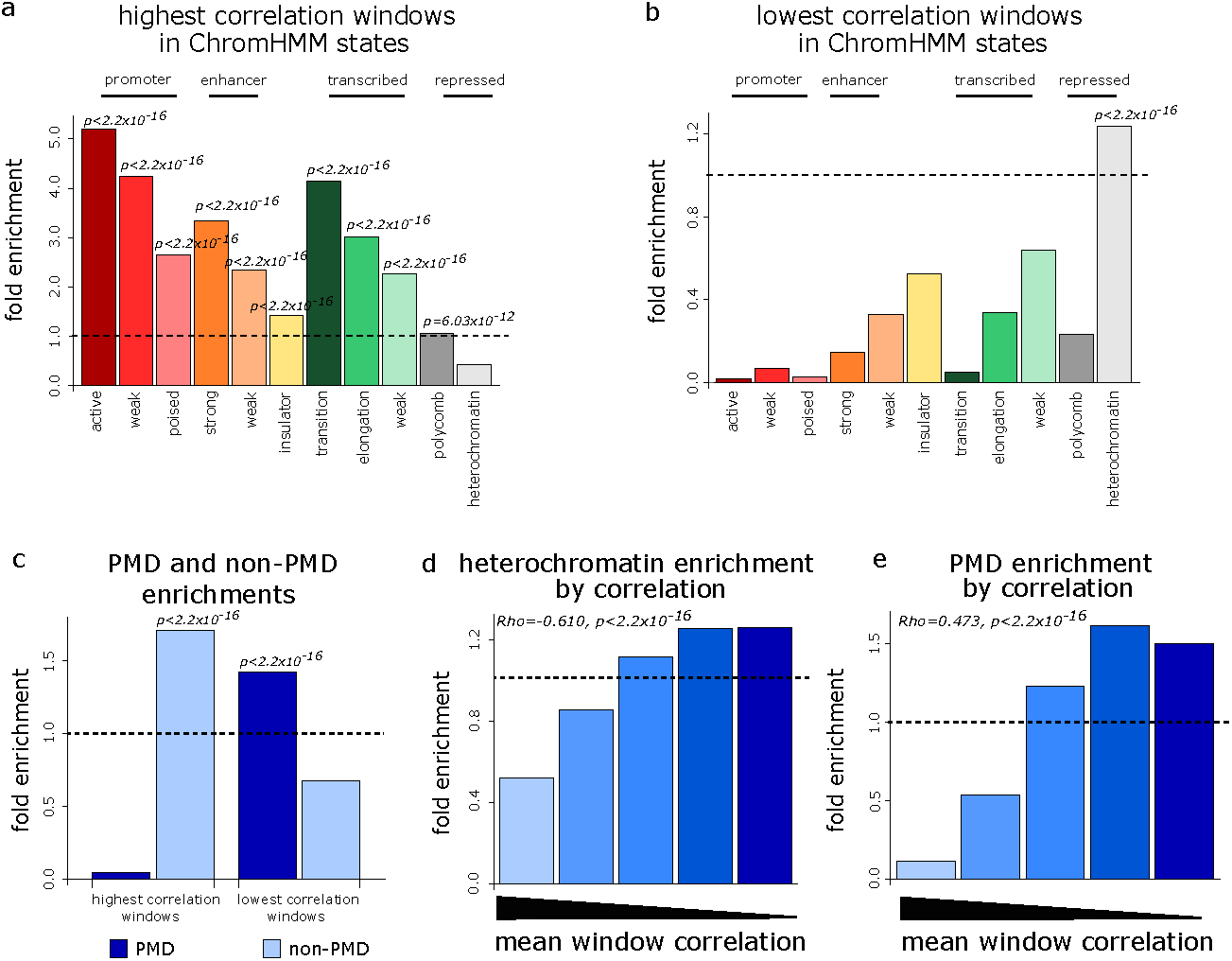
Heterochromatin possesses heterogeneous methylation patterns. **a**,**b** Barplots showing enrichment of the 10% least (**a**) and most (**b**) heterogeneous 100kb windows in chromHMM states defined in GM12878 lympoblastoid cells. **c** Barplot showing the enrichment of the 10% least and most heterogeneous 100kb windows in PMDs and non-PMDs. **d** Barplot showing enrichment of continuous heterogeneity groups of genomic windows (defined from the ranked mean correlation) in the chromHMM heterochromatin annotation from GM12878 lymphoblastoid cells. **e** Barplot showing enrichment of continuous heterogeneity groups of genomic windows (defined from the ranked mean correlation) in PMDs. In **d, e** the Spearman correlations (Rho) between mean window correlation and window overlap with heterochromatin or PMDs, respectively, are also shown along with the relevant p-values.

In some somatic cells, heterochromatin is associated with regions of intermediate methylation levels that are termed partially methylated domains (PMDs) [3, 24]. To further assess the correspondence between heterochromatin and intra-molecular DNA methylation heterogeneity, we therefore called 678 PMDs in GM24385 cells from our Nanopore DNA methylation data (Fig. S4**c**) and defined the rest of the genome as 582 non-PMDs. GM24385 PMDs showed substantial overlap with those previously defined in 11 other lymphoblastoid cells from WGBS data (Jaccard = 0.804) [25]. Consistent with their enrichment in the chromHMM-defined heterochromatin state, genomic windows with the lowest mean correlation were significantly enriched in GM24385 PMDs (Fig 4**c**, Wilcoxon test *p <* 2.20 × 10^*−*16^) with 60.0% of these windows overlapping PMDs by at least half the window length. Those windows with the highest mean correlation were significantly enriched in non-PMDs (Fig 4**c**, Wilcoxon test *p <* 2.20 × 10^*−*16^) with 97.6% of these windows overlapping non-PMDs by at least half the window length. We observed similar enrichment results when the analysis was repeated using 50kb and 200kb genomic windows (Fig. S4**d**) suggesting that overlap between intra-molecular methylation heterogeneity and PMDs was independent of the choice of genomic window size.

To examine how methylation heterogeneity associates with heterochromatin and PMDs more generally, we calculated the correlation between the mean correlation in 100kb genomic windows and the proportion of window overlapping the chromHMM heterochromatin state or PMDs. Since methylation heterogeneity increases as correlation decreases, a negative correlation between correlation and overlap with a genomic annotation is indicative of the annotation being associated with high heterogeneity. Mean window correlation showed a significant negative correlation with both the proportion of the window covered by the chromHMM heterochromatin state and the proportion of the window covered by PMDs (Fig 4**d,e**, Spearman correlation = − 0.610 and − 0.473 respectively, both *p <* 2.20 × 10^*−*16^). At the single-molecule level, we also found significant negative correlations between a read’s correlation and the proportion of the read that overlapped the heterochromatic state and PMDs (Spearman correlation = − 0.474 and − 0.351 respectively, both *p <* 2.20 × 10^*−*16^) confirming that the association between intra-molecular DNA methylation heterogeneity and heterochromatin was maintained at the single-read level.

Taken together, these analyses suggest that the regions of the human genome displaying the greatest degree of large-scale intra-molecular methylation heterogeneity are heterochromatic partially methylated domains.

### Periodic DNA methylation patterns are observed in single reads with high heterogeneity

Having shown that heterogeneity in DNA methylation patterns is observed in heterochromatin, we now sought to determine the nature of intra-molecular DNA methylation heterogeneity in these regions in order to understand its potential causes.

In order to do so, we first defined groups of single reads exhibiting low or high intra-molecular methylation heterogeneity by fitting the sum of two Gaussian distributions to the bimodal distribution of correlations (Fig 2**a**: low correlation reads, i.e. high heterogeneity reads, and high correlation reads, i.e. low heterogeneity reads, correspond to the left and right peaks respectively). High heterogeneity reads were significantly enriched in heterochromatin and PMDs, while low heterogeneity reads were enriched in promoters, enhancers, insulators, transcribed regions, polycomb-repressed regions and non-PMDs, confirming the associations observed in previous analyses (Wilcoxon tests, *p <* 2.20 × 10^*−*16^ in each case).

We now sought to determine how the correlation between CpGs depends on the distance between them. Analysis of bulk WGBS data has reported oscillatory distance-dependent correlations in mean methylation levels within PMDs [26–28]. Given that we observed high intra-molecular heterogeneity associated with PMDs, we wondered whether oscillatory DNA methylation patterns might be present at the single-molecule level and account for some of the heterogeneity we measure. We therefore calculated the correlation in methylation state for pairs of CpGs separated by different distances in single reads. Calculating these correlations for all possible distances in each individual long read is prohibitively computationally expensive. To reduce the computational burden, we therefore calculated distance-dependent correlations within low and high heterogeneity reads for distances up to a maximum of 500bp. As expected, mean distance-dependent correlations were higher for low heterogeneity reads than high heterogeneity reads across distances of 2-500bp (Fig 5**a**). Within both low and high heterogeneity reads, the mean distance-dependent correlation was highest for CpGs located close to each other and reduced as the distance between CpGs increased (Fig 5**a**). However, after initially falling, the mean distance-dependent correlation for high heterogeneity reads then increased, showing oscillatory behaviour with a wavelength of around 185bp (Fig 5**a**). No clear oscillatory behaviour was apparent in the mean distance-dependent correlation profile for low heterogeneity reads (Fig 5**a**). To determine whether the oscillatory behaviour within high heterogeneity reads could be seen beyond distances of 500bp, we then computed distance-dependent correlations up to 2kb between CpGs within these reads. While the oscillations decrease in magnitude with distance (Fig 5**b**), we observed that they continued up to 2kb within high heterogeneity reads (Fig 5**c**). Since we previously found PMDs to be associated with highly heterogeneous single-molecule methylation patterns, we asked whether oscillatory patterns were also visible in reads aligning to PMDs. Indeed, the distance-dependent correlations calculated for PMD reads showed oscillatory behaviour up to 2kb similarly to the high heterogeneity reads (Fig. S5**a**,**b**).

**Fig 5.**
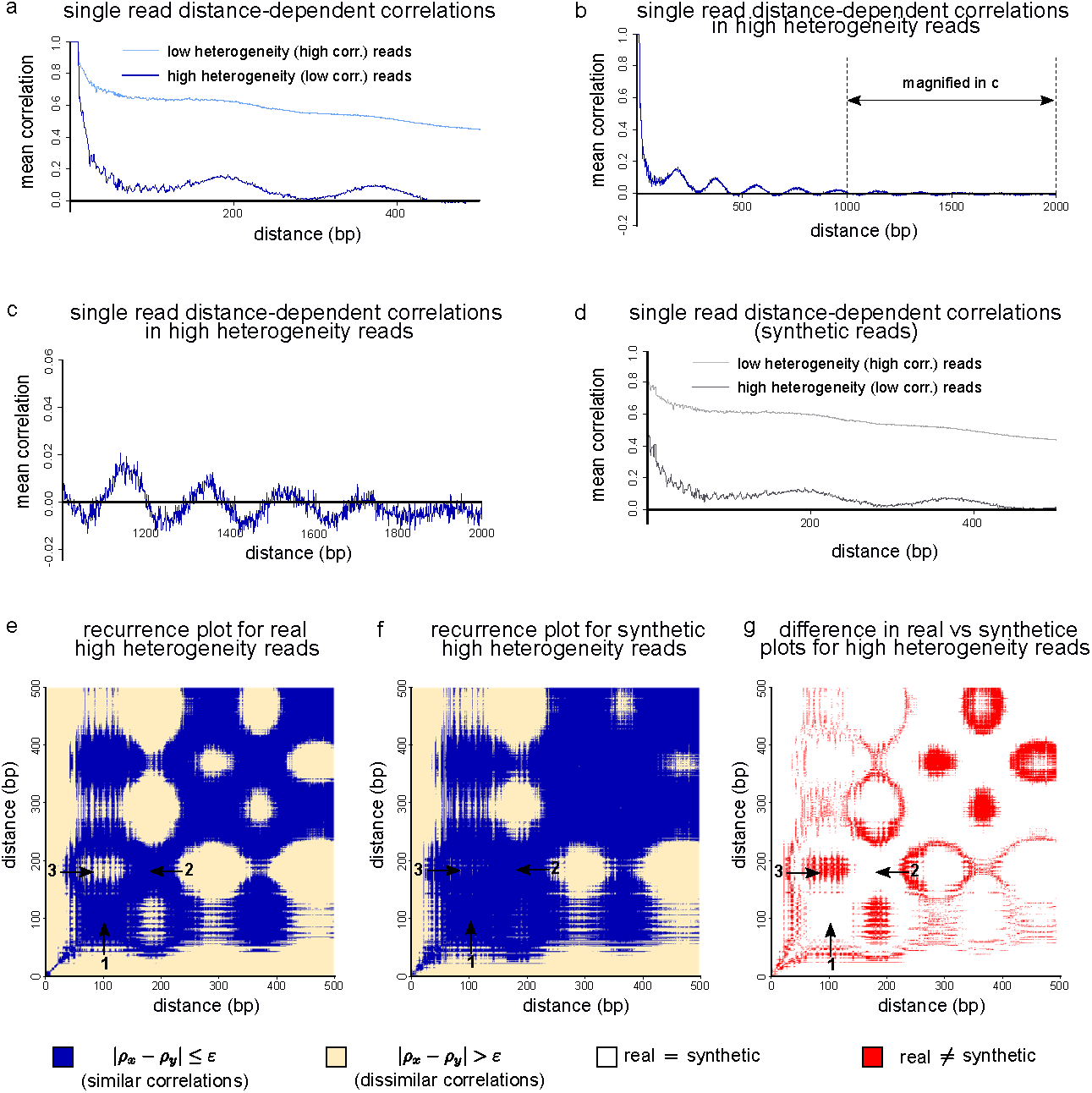
Periodic DNA methylation patterns are observed in single molecules with a high heterogeneity. **a** Plot of mean single-read distance-dependent correlation between CpG sites within 500bp of each other for 1, 461, 561 high heterogeneity reads and 544, 193 low heterogeneity reads. **b** Plot of mean single-read distance-dependent correlation between CpG sites within 2kb of each other for 1, 461, 561 high heterogeneity reads. **c** Magnification of the region indicated in **b** (i.e. the mean single-read distance-dependent correlation for distances 1kb-2kb in high heterogeneity reads). **d** As in **a** but for synthetic reads generated based on bulk methylation properties. **e**,**f** Recurrence plots for real (**e**) and synthetic (**f**) high heterogeneity reads for CpGs within 500bp of each other (see Methods). For these plots, *ε* = 0.06. Arrows 1 and 2 indicate example regions where the correlations for distances *x* and *y* are similar in the real reads. Arrow 3 indicates an example region where the correlations for distances *x* and *y* are dissimilar in the real reads. **g** Difference plot derived from the recurrence plots in (e) and (f) showing agreement between the real and synthetic data in white and disagreement in red. The regions indicated by arrows 1 and 2 show agreement between the real and synthetic data. However the real and synthetic data disagree in the region indicated by arrow 3.

We now aimed to determine the degree to which the single-molecule methylation patterns we observed were a consequence of the bulk mean methylation levels. Since the methylation state of each CpG in the single-molecule data is described using a binary variable, while the bulk mean methylation level of each CpG is a continuous variable, it is not possible to directly compare single-molecule states with bulk mean methylation levels. We therefore generated synthetic single-molecule data based on the bulk methylation data. In particular, for each read, we generated a synthetic read where the methylation state of each CpG in the synthetic read is determined by the probability that it is methylated in the bulk data (see Methods). We then calculated distance-dependent correlations up to 500bp from these synthetic reads and compared them to what we observed in the real reads. The mean distance-dependent correlations derived from synthetic high heterogeneity reads showed oscillatory behaviour; however, the magnitude of the oscillations was lower than that observed from the real high heterogeneity reads (Fig 5**d**). As in the real low heterogeneity reads, no clear oscillatory behaviour was observed in the synthetic low heterogeneity reads (Fig 5**d**).

These analyses suggested that the patterns within single high heterogeneity reads differed from that expected from mean bulk methylation levels. To further understand these differences, we used recurrence plots to examine distance-dependent correlations calculated up to 500bp for real and synthetic single reads. Recurrence plots use a binary colouring system which makes patterns within spatial and temporal data easily observable [29–31]. A recurrence plot is composed of a grid and, in the context presented here, the colour of block (*x, y*) in the grid is one of two colours depending on whether the correlations associated with distances *x* and *y* are within a pre-specified tolerance of each other. The recurrence plot associated with the real high heterogeneity reads showed alternating areas of high and low similarity Fig 5**e**. In particular, we observed large, regular blue areas corresponding to regions of high similarity in the correlations associated with different distances *x* and *y* (Fig 5**e** arrows 1 and 2, respectively). These had distinct boundaries from yellow elliptical regions corresponding to regions of low similarity (Fig 5**e** arrow 3). The recurrence plot associated with the synthetic high heterogeneity reads was similar. However, the observed structure in methylation patterns was less distinct than that of the real high heterogeneity reads, with correlations being generally more similar across distances (Fig 5**f**). Differences between the recurrence plots for the real and synthetic reads were concentrated at the boundaries separating regions where the correlations are similar from regions where the correlations are dissimilar (Fig 5**g** red regions, e.g. arrow 3). These analyses demonstrated that the structured single-molecule methylation patterns associated with heterochromatin and PMDs differ from those predicted from the bulk mean methylation levels.

The ∼ 185bp wavelength of the oscillations observed in the high heterogeneity reads lies within the range of reported 160 − 230bp nucleosomal DNA repeat lengths [32]. It has also been postulated that nucleosomes underpin oscillatory patterns observed in PMD bulk mean DNA methylation levels [26–28]. At the single-molecule level, the length of linker DNA between nucleosomes can vary and nucleosomes can be differentially phased (positioned) between molecules [33]. To gain intuition as to whether nucleosome organisation at the single-molecule level could generate the differences we observed between the real reads and synthetic reads generated from bulk data, we considered simple proof-of-principle models of extreme cases of nucleosome placement and interaction with DNA methylation on single molecules.

To do so, we assumed that each nucleosome occupied 147bp of DNA, linker length varied between 10bp and 90bp, and methylation state was entirely dictated by nucleosome occupancy (see Methods). We first generated single-molecule methylation data using a model where nucleosomes were identically spaced and phased between molecules (Fig. S5**c**, see Methods) before calculating single-molecule distance-dependent correlations from this model and for modelled synthetic reads derived from its bulk properties. The single-molecule recurrence plot derived from the distance-dependent correlations qualitatively captured the patterns we observed in the real high heterogeneity reads (Fig. S5**d** vs. Fig 5**e**). This suggests that nucleosomes could play a role in generating the heterogeneous DNA methylation patterns we see in single molecules. In this case of perfect phasing between molecules, we also observed that the recurrence plot generated using the bulk properties of the model was identical to that of the modelled single-molecule data (Fig. S5**d, e**).

We next generated data from a model where nucleosome positioning was completely unphased between molecules (Fig. S5**f**, see Methods). Again, the single-molecule recurrence plot derived from the model qualitatively captured the true patterning observed for the real high heterogeneity reads (Fig. S5**g** vs. Fig 5**e**). However, the recurrence plot generated using the bulk properties of the model showed less distinct boundaries between regions corresponding to similar and dissimilar correlations than was observed in the single-molecule recurrence plot generated from the model (Fig. S5**g, h**). This is in qualitative agreement with the differences we observed in the recurrence plots generated from real and bulk-simulated high-heterogeneity reads (Fig 5**e—g**) consistent with the hypothesis that these differences could be generated by differences in nucleosomal organisation between molecules.

These analyses demonstrate that the parts of the human genome showing high intra-molecular DNA methylation heterogeneity display kilobase-scale oscillatory DNA methylation patterns within single molecules. These patterns differ from those predicted from mean bulk DNA methylation levels and could result from differences in nucleosome phasing between molecules.

## Discussion

Here, we have used Nanopore sequencing data to analyse large-scale intra-molecular DNA methylation heterogeneity in the human genome. Our results show that intra-molecular heterogeneity in DNA methylation patterns is abundant and underestimated by bulk-level metrics. We specifically observe a high degree of intra-molecular DNA methylation heterogeneity in heterochromatic regions and that this heterogeneity manifests as oscillatory DNA methylation patterns that differ from those predicted from mean bulk methylation levels (Fig 6**a**).

**Fig 6.**
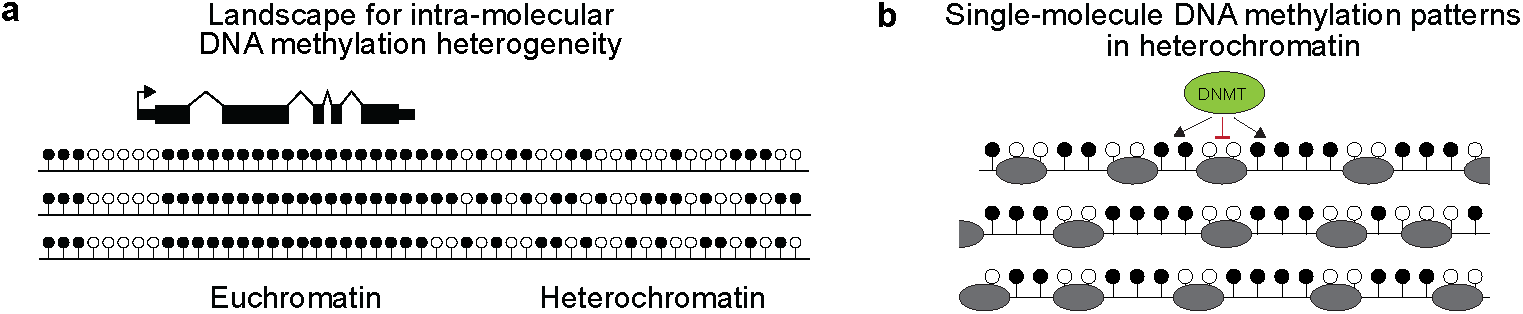
Intra-molecular DNA methylation heterogeneity varies across the human genome and is affected by nucleosomes at the single-molecule scale. **a** Heterogeneous DNA methylation patterns are observed in heterochromatin. Schematic model of the arrangement of intra-molecular DNA methylation heterogeneity in the genome. Euchromatic, genic regions possess ordered, uniform patterns whereas heterochromatin is characterised by more heterogeneous patterns at the single-molecule level. **b** Oscillatory DNA methylation patterns are observed in single molecules within heterochromatin and could be caused by differences in nucleosomal organisation between molecules. Schematic showing proposed arrangement of DNA methylation and nucleosomes in single DNA molecules in heterochromatin. We propose that periodic patterns result from methylation of linker DNA due to nucleosomes being refractory to DNMTs [36–39]. Differences in phasing and positioning between molecules cause sites of methylation to be imperfectly aligned across molecules meaning that these patterns are partially masked in bulk DNA methylation data.

We propose that nucleosomes are a likely source of the heterogeneity we observe in heterochromatin. The total length of nucleosomal and inter-nucleosomal linker DNA varies in the range of 160-230bp depending on the organism, cell type and region of the genome examined [32–35]. *In vitro* experiments demonstrate that the wrapping of DNA around nucleosomes is refractory to methylation by DNMTs [36–39] so that methylation occurs at the linker DNA [40]. This suggests that the blocking of DNMTs by nucleosomes could generate the oscillations of around 185bp that we observe in reads with high intra-molecular heterogeneity (Fig 6**b**). *In vivo* linker DNA methylation is seen in between the well-positioned nucleosomes around CTCF sites in human cells [41], consistent with the hypothesis that nucleosomes are refractory to DNMTs. A similar organisation has also been reported at CTCF sites at the single-molecule level [42].

However, only an estimated 8.7% of nucleosomes in the human genome are strongly-positioned [43] and the majority are not phased between molecules [33]. A single-molecule analysis of Micrococcal nuclease digested fragments suggests that heterochromatin is characterised by regularly spaced nucleosomal arrays and that spacing is more irregular in euchromatin [44]. Regular nucleosome spacing in heterochromatin was also observed in a single-cell nucleosomal positioning study which additionally suggested that heterochromatin is characterised by a lack of phasing between molecules [45]. In addition, a single-molecule footprinting analysis of oligonucleosomes released from cells found that irregular oligonucleosome patterns were abundant in heterochromatin [46]. However, despite the reported lack of phasing of heterochromatic nucleosomes, analyses of bulk mean DNA methylation levels has reported structured periodicity in DNA methylation patterns in PMDs [26–28]. Here we show for the first time that periodicity in heterochromatic DNA methylation patterns manifests at the kilobase scale within single molecules. We also find that the patterns we observe at the single-molecule level are not completely predicted from mean bulk DNA methylation levels; in fact they are partially masked. Our simple proof-of-principle models are consistent with the hypothesis that this masking could result from differences in the phasing and organisation of nucleosomes between molecules (Fig 6**b**). We note that the oscillations previously observed in the distance-dependent correlations from heterochromatic bulk data could suggest heterochromatic nucleosomes show some degree of phasing and preferential positioning rather than being entirely unphased between molecules. Oscillations were not observed in the low heterogeneity reads associated with euchromatic, genic regions. We suggest that this could occur because nucleosome remodelling, for example through the action of FACT in displacing nucleosomes during transcription [47], could enable DNMTs to access nucleosomal DNA.

Previous work has also suggested that DNA sequence plays a strong role in shaping bulk mean DNA methylation patterns in PMDs. The sequence immediately flanking a CpG, as well as local CpG density are associated with, and can predict, PMD bulk mean methylation levels [26]. CpGs lacking neighbours and flanked by A or T bases have particularly low bulk mean methylation levels [48]. Such CpGs also lose methylation with successive cell divisions *in vitro* [49] and have been reported to have slower remethylation kinetics [50]. Deep-learning models can also accurately predict methylation loss based on the sequence of the 150bp surrounding a CpG [51]. However, nucleosome positioning is also strongly influenced by sequence [52] and DNMT activity varies depending on CpG flanking sequences [53]. It is therefore possible that observations of the influence of sequence and nucleosomes on DNA methylation in heterochromatin are linked and future studies will be required to precisely dissect their respective roles.

A variety of statistics have previously been used to examine heterogeneity in DNA methylation patterns. These include the proportion of discordant reads (PDR) [54], methylation haplotype load (MHL) [55] and fraction of discordant reads (FDR) [19]. However, these measures quantify inter-molecular heterogeneity and so are not directly comparable to our analysis of intra-molecular heterogeneity. An information theory approach has been used to predict probability distributions associated with single-molecule methylation patterns from WGBS data and suggested that the genome is organised into domains of disordered and ordered methylation patterns that coincide with large-scale chromatin organisation [10]. However, although these studies are consistent with our observation of high intra-molecular heterogeneity in heterochromatin, WGBS reads are not sufficiently long to directly assess single-molecule DNA methylation patterns across the scale of 10^4^-10^5^ bases as we have done here.

## Conclusion

Taken together our observations suggest that the degree of intra-molecular heterogeneity in DNA methylation patterns varies widely across the human genome. We propose that the nature of this heterogeneity suggests a role for nucleosomes in shaping DNA methylation patterns at the single-molecule level, particularly in heterochromatin.

## Materials and methods

Unless otherwise specified, we performed analysis using R (v. 4.1.2), Mathematica (v. 12.3) and Bedtools (v. 2.30.0) [56]. All statistical tests were two-sided.

### Preprocessing of Nanopore data

We downloaded raw Nanopore reads in fast5 format for GM24385 lymphoblastoid cells from Oxford Nanopore (amazon bucket s3://ont-open-data/gm24385_2020_09/). We then basecalled these reads using the dna_r9.4.1_450bps_hac_prom.cfg configuration of Guppy (v. 5.0.11) [57], aligned them to the human genome (hg38 assembly) using the map-ont setting of Minimap2 (v. 2.22) [58] and called the methylation state at CpGs using Nanopolish with default settings (v. 0.13.2) [18]. Methylation states were assigned to each CpG site based on the log likelihood ratio (LLR) calculated by Nanopolish. We used the LLR cut-off used by Nanopolish scripts and estimated to have an accuracy *>* 92.5% [18]. Comparisons to WGBS also suggest Nanopolish methylation calls are accurate [59, 60]. Here we consider single-molecule patterns ≥ 100 CpGs in length so the influence of a small number of miscalled methylation states is likely to be minimal. CpG sites were assigned an unmethylated status if LLR ≤ − 2 and a methylated status if LLR ≥ 2. For |LLR| *<* 2 we deemed the CpG site to be unassignable. Using these settings, 72.6% of observations were assigned methylated or unmethylated states. Only reads mapping to canonical hg38 autosomes (chromosomes 1-22) were considered in our analyses.

### Calculation of single-read statistics

We converted methylation tsv files outputted by Nanopolish to bed format using the mtsv2bedGraph.py script from the timplab/nanopore-methylation-utilities GitHub repository [42]. We used a custom R script to identify the 2, 908, 181 out of 6, 289, 480 reads that had defined methylation states for ≥ 100 CpGs. For each retained read, *r*, we discarded CpG sites that could not be assigned a methylation state and stored the methylation states of the remaining *N*_*r*_ ≥ 100 CpGs in a vector 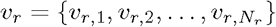, where for *i* = 1, … *N*_*r*_,

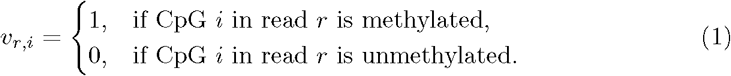

For each read *r* we then used R (v. 4.0.1) to calculate the mean methylation level (*μ*_*r*_), coefficient of variation (CV_*r*_) and the correlation in methylation level between neighbouring sites (*ρ*_*r*_) via

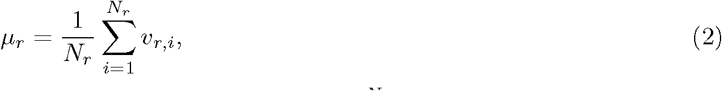

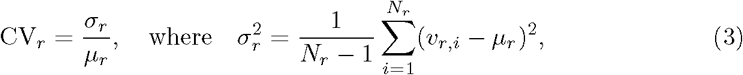

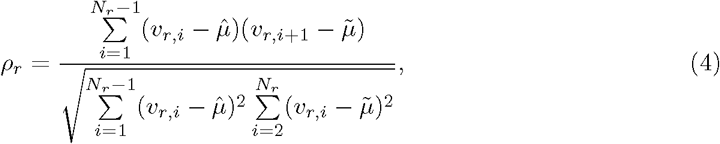

where

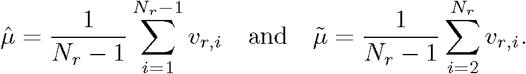

### Definition of genomic windows

We split canonical autosomal chromosomes into 28,760 non-overlapping genomic windows of 100kb using BEDOPS (v. 2.4.26) [61]. Note that 22 windows (corresponding to the end of each autosome) were shorter than 100kb. This process was repeated but with 50kb and 200kb windows for selected analyses.

### Calculation of mean single-read statistics

We used Bedtools and custom command line scripts to extract the 1, 707, 118 reads that were entirely contained within a single window and had called methylation states for ≥ 100 CpGs. We then calculated mean single-read statistics associated with each window using R (v. 4.0.1). In particular, if 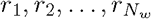 were the reads that aligned entirely within a window, *n*, then we calculated the single-molecule mean (*μ*_*s,n*_), coefficient of variation (CV_*s,n*_) and correlation (*ρ*_*s,n*_) associated with window *n* via

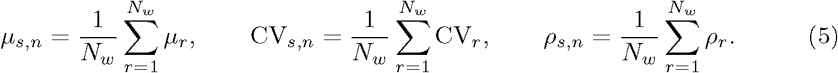

This lead to mean single-molecule statistics being obtained for 26, 951 100kb windows.

### Calculation of bulk-level window statistics

We defined the methylation coverage for each CpG site to be the combined number of methylated and unmethylated calls associated with that CpG site using all of the 6, 289, 480 reads that covered that CpG. We discarded CpG sites with *<* 10 methylation calls and those that overlapped the boundary between two windows (i.e. if the forward-strand cytosine belongs to one window and the guanine to another). The mean CpG methylation for each site was defined as the number of methylated calls associated with that CpG divided by the total number of calls (methylated and unmethylated) observed for that CpG. We then removed 100kb windows with *<* 100 retained CpG sites. We used the CpG methylation levels associated with the remaining 26, 714 windows to calculate bulk-level statistics for each window in R (v. 4.0.1). In particular, for window *n* with *N* ≥ 100 retained CpGs with methylation levels *m*_1_, *m*_2_, …, *m*_*N*_, we calculated the bulk mean methylation level (*μ*_*b,n*_), coefficient of variation (CV_*b,n*_) and the correlation in methylation level between nearest-neighbour sites (*ρ*_*b,n*_) via

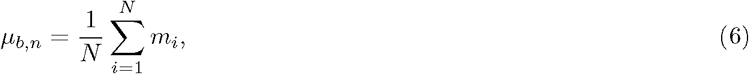

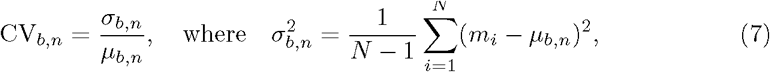

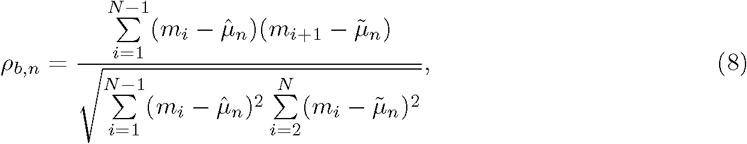

where

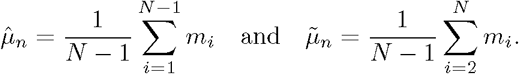

### Comparison of mean single-molecule and bulk-level window statistics

We used Spearman correlation tests in R to calculate correlations and associated p-values between the bulk statistics and mean single-molecule statistics for the 26, 687 100kb genomic windows that had sufficient bulk and single-molecule data. We also fit linear models to the bulk-level and single-molecule statistics using the base lm command in R and conducted paired t-tests to assess the difference between the bulk and single-molecule values. The average absolute relative difference between the bulk and single-molecule mean was calculated via

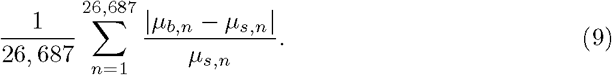

Similarly, the average absolute relative difference between the bulk and single-molecule coefficient of variation and correlation were calculated via

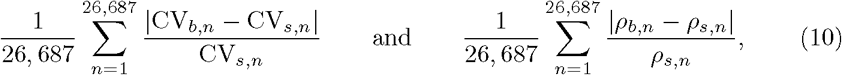

respectively.

### Generation of simulated and shuffled correlations from individual reads

We simulated the case where methylation states were randomly distributed along each read. For each read *r* with methylation states for ≥ 100 CpGs, we created a simulated read containing the same number of CpGs, but where the methylation state of each CpG was randomly chosen from a binomial distribution with success rate for methylation equal to the mean methylation level of the read (*μ*_*r*_). The correlation was then calculated for each of these synthetic reads. The correlation distribution obtained from the true reads was compared to that obtained from the synthetic reads and a Kolmogorov-Smirnov test was conducted in R to assess whether the observed methylation patterns in true reads are random or non-random.

To generate the mean shuffled correlation for genomic windows we calculated a mean correlation for each window using random reads (rather than those that align to that window). Reads were shuffled using the command-line *shuf* command. For example, if 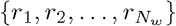 were the reads aligning entirely within window *n*, then we replaced these by a set of random reads 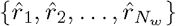, and calculated a mean correlation for window *n* via

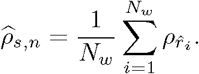

We then compared the *ρ*_*s,n*_ and 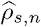 distribution using a Kolmogorov-Smirnov test in R.

### Calculation of RTS

For each read *r* containing methylation states for ≥ 100 CpGs, we calculated the read transition score in R (v. 4.0.1) as

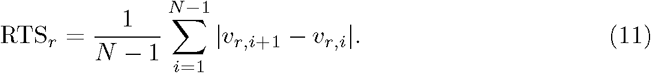

RTS_*r*_ essentially measures the probability that two nearest-neighbour CpG sites in read *r* have different methylation states. Reads which are entirely unmethylated or entirely methylated therefore have RTS 0, while reads where no two nearest-neighbour CpG sites share the same methylation state have RTS 1.

Mean RTSs were then calculated for each genomic window similarly as for the other mean single-read statistics. In particular, if 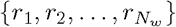 were the reads that aligned entirely within a window, *n*, then we took the mean RTS associated with window *n* to be

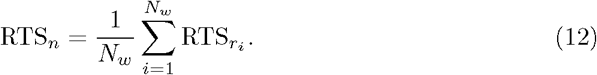

Shuffled mean RTSs were also obtained for each window in a similar manner (see Methods).

### Defining windows with the lowest and highest intra-molecular heterogeneity

We ranked the 26, 951 windows by *ρ*_*s,n*_ before defining the windows with the greatest intra-molecular heterogeneity as the 10% of windows with the lowest mean correlation and the least heterogeneous windows as the 10% of windows with the highest mean correlation.

### Comparison of intra-molecular heterogeneity levels to inter-molecular heterogeneity levels

We computed the measure *d*_*i*_ = |*m*_*i*_ − 0.5|, where *m*_*i*_ is the methylation level of CpG *i* for each CpG with ≥ 10 methylation calls. *d*_*i*_ = 0.5 indicates low molecule-to-molecule heterogeneity for CpG *i*, where as *d*_*i*_ = 0 corresponds to high molecule-to-molecule heterogeneity for CpG *i*. We performed Kolmogorov-Smirnov tests to compare the distribution of *d*_*i*_ associated with the 10% of windows with the lowest mean correlation and the distribution associated with the rest of the genome and to compare the 10% of windows with the highest mean correlation and the distribution associated with the rest of the genome. In particular, we used Kolmogorov-Smirnov tests in R to test whether the cumulative distribution function (CDF) associated with *d*_*i*_ for the lowest or highest mean correlation windows is significantly different than that associated with the rest of the genome.

### Sourcing and processing of genomic annotations

Using the command line, we derived bed files of hg38 gene locations using annotation files from Gencode (v. 41) [62]. CGI annotations were taken from [63]. Overlapping CGI intervals were merged using BEDtools (v. 2.27.1) before they were converted to hg38 positions using the UCSC browser liftover tool (https://genome.ucsc.edu/cgi-bin/hgLiftOver). Non-autosomal CGIs were excluded from the analysis as were CGIs located in ENCODE blacklist regions [22] as previously described [64].

We downloaded hg19 GM12878 lymphoblastoid cell chromHMM states from UCSC (http://www.genome.ucsc.edu/cgi-bin/hgTables), [22, 23] before lifting them over to hg38 genome coordinates (https://genome.ucsc.edu/cgi-bin/hgLiftOver). To remove cases where multiple hg19 coordinates mapped to the same hg38 coordinate, we used the command line and Bedtools to remove from the analysis any regions associated with more than one annotation type in hg38 as a consequence of this liftover.

We downloaded PMD locations identified for 11 lymphoblastoid cell lines from the AllPMDs.tar.bz2 file in the bdecatoPMD Paper Scripts GitHub repository [25]. These coordinates were then converted from hg19 to the hg38 coordinates using Liftover (https://genome.ucsc.edu/cgi-bin/hgLiftOver). We excluded poorly mapped regions of the genome using annotations of gaps and centromeres from the UCSC browser (hg38 gap and centromere tracks). Annotations were downloaded from the UCSC table browser. Regions annotated as heterochromatin, short arm and telomeres from the gaps track were merged with the centromeres track using Bedtools merge with -d set to 10Mb. This merged file was then excluded from the PMD BED files using Bedtools subtract.

### Definition and analysis of PMDs

We called PMDs for GM24385 cells using Methpipe (v. 5.0.0) [25] on mean bulk CpG methylation levels and CpG methylation coverage calculated from Nanopore data. We defined non-PMDs as genomic regions that did not overlap with a PMD. Poorly mapped regions of the genome were removed from the PMD and non-PMD annotations as previously discussed. Using Bedtools we also excluded PMDs and non-PMDs shorter than 200kb. We compared the overlap between GM24385 PMDs to those from other lymphoblastoid cells [25] using the Bedtools Jaccard function on a BED file generated by merging the PMDs annotated in the other cell lines using Bedtools merge.

### Testing enrichment of annotations in genomic windows

For each annotation type, *a*, we used Bedtools to compute

- *P*_*a*,all_ : the proportion of bases in all 26, 951 windows analysed that overlap the annotation (i.e. the sum of bases from all windows that overlap the annotation divided by the summed length of all windows);
- *P*_*a*,least_ : the proportion of bases in the least heterogeneous windows that overlap the annotation (i.e. the sum of bases from the least heterogeneous windows that overlap the annotation divided by the summed length of the least heterogeneous windows);
- *P*_*a*,most_ : the proportion of bases in the most heterogeneous windows that overlap the annotation (i.e. the sum of bases from the most heterogeneous windows that overlap the annotation divided by the summed length of the most heterogeneous windows).

We then calculated the enrichment of the least and most heterogeneous windows in annotation type *a* using

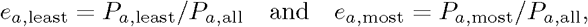

respectively. To test the significance of the enrichment, we extracted the proportion that each window overlaps annotation type *a* using Bedtools and compared the distribution of proportions associated with the least/most heterogeneous windows to the proportions associated with all windows using Wilcoxon tests in R.

### Testing continuous relationship between correlation and annotations

To test the continuous relationship between a windows mean correlation and the proportion of overlap with genomic annotations, we calculate the correlation between a window’s mean correlation and its overlap with heterochromatin and PMDs. For graphing, we split windows into five equally sized groups based on their ranked mean correlation. We then repeated the enrichment analyses previously described to calculate the annotation enrichment level in each group.

To test the relationship between correlations within single reads and overlap with genomic annotations, we used Bedtools to extract the proportion of each read that overlapped the chromHMM heterochromatin state or PMDs and conducted Spearman correlation tests in R to compare these overlaps to the read correlations.

### Derivation of low and high correlation reads by mixture modelling

Using Mathematica we fit the sum of two Gaussian distributions, with means *μ*_1_, *μ*_2_ (*μ*_1_ *< μ*_2_) and standard deviations *σ*_1_, *σ*_2_, to the bimodal distribution for the read correlation (Fig 2**a**). Of the 2, 907, 431 reads containing methylation information for a minimum of 100 CpG sites and for which the correlation between nearest-neighbour sites could be computed, we defined the 1, 461, 561 reads that satisfied |*ρ*_*r*_ − *μ*_1_| ≤ *σ*_1_ to be the “low correlation group” (or “high heterogeneity group”) and the 544, 193 reads that satisfied |*ρ*_*r*_ − *μ*_2_| ≤ *σ*_2_ to be the “high correlation group” (or “low-heterogeneity group”), where *ρ*_*r*_ is the correlation associated with read *r*. We discarded the remaining reads. The previous enrichment analyses were then repeated for the low and high correlation read groups.

### Calculating distance-dependent correlations

For each read in the high heterogeneity (i.e. low correlation) group, we extracted the methylation state of each possible pair of CpGs along with the distances between those CpGs. In particular, for a read *r* covering *N*_*r*_ CpG sites, we extracted {*u*_*i*_, *u*_*j*_, *d*_*i,j*_} for each *i* ∈ {1, 2, …, *N*_*r*_ − 1}, *j* ∈ {*i* + 1, …, *N*_*r*_}, where

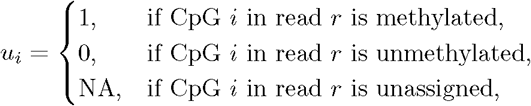

and *d*_*i,j*_ is the distances (in bps) between CpG *i* and CpG *j* in read *r*. For each distance *d* = {2, 3, …, 2000} we extracted {*u*_*i*_, *u*_*j*_} corresponding to *d*_*i,j*_ = *d, u*_*i*_ ≠ NA and *u*_*j*_ ≠ NA to obtain a matrix of states *S* with *L* ≥ 0 rows, where each row denotes the methylation state of a pair of CpGs lying a distance of *d* apart from each other in read *r*. If *L* ≥ 2, then we used R (v. 4.0.1) to compute the correlation between sites on read *r* a distance of *d* apart via

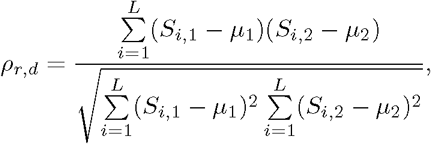

where

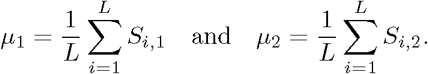

For each *d* = {2, 3, …, 2000} we computed the mean of *ρ*_*r,d*_ over all reads in the high heterogeneity group to obtain mean single-molecule distance-dependent correlations associated with the high heterogeneity group.

We obtained mean single-molecule distance-dependent correlations associated with the low heterogeneity group analogously for distances 2-500bp.

Reads aligning entirely within PMDs were also extracted using Bedtools and mean single-molecule distance-dependent correlations associated with these reads were calculated for distances 2-2000bp as above.

### Generation of synthetic reads from bulk DNA methylation data

We edited the calculate methylation frequency.py script provided by Nanopolish [18] to output the proportion of times that each CpG site was called to be unmethylated, methylated or unassigned across reads, where again each CpG site was called to be methylated if LLR ≥ 2, unmethylated if |LLR| ≤ − 2 and unassigned if LLR *<* 2. For each read in the Nanopore dataset we then generated a synthetic read based on this bulk information. Explicitly, for each read *r*, a synthetic read 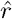 was created where the methylation state of CpG *i* in read 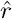 is set to be unmethylated with probability *μ*_*u,i*_, methylated with probability *μ*_*m,i*_ and unassigned with probability *μ*_NA,*i*_, where *μ*_*u,i*_, *μ*_*m,i*_, *μ*_NA,*i*_ are the bulk probabilities that CpG *i* is unmethylated, methylated or unassigned.

The synthetic reads corresponding to the low and high heterogeneity reads were then extracted and used to calculated distance-dependent correlations as previously.

### Generation of recurrence plots

Recurrence plots were generated using the Mathematica ListDensityPlot command. In the recurrence plots, block (*x, y*) was set to be blue if |*ρ*_*x*_ − *ρ*_*y*_| ≤ *ε* for some *ε >* 0, where *ρ*_*x*_ and *ρ*_*y*_ are the correlations associated with distances *x* and *y*, respectively, and is yellow otherwise. We chose *ε* = 0.06 as this was the tolerance value for which the Euclidean distance between the real and synthetic recurrence plots is maximised. Recurrence plots were obtained for a range of *ε* to check consistency of observations (data not shown).

### Mathematical models of nucleosome occupancy

We generated simple models of the interaction between DNA methylation and nucleosomes using Mathematica. As nucleosomes are refractory to the action of DNMTs, we assumed that nucleosome occupied DNA was deterministically unmethylated and linker DNA was methylated [36–40]. We also assumed that nucleosomes occupied 147bp of DNA and that linker length varied uniformly within the range 10 to 90bp [32]. To simplify the models further and make them more tractable, we assumed that every DNA base on the molecules could be methylated. Motivated by this, we simulated two mathematical models of nucleosome occupancy. In each model, nucleosome occupancy data was simulated for *m* = 200 reads (*R*_1_, …, *R*_200_) covering the same region of length *n* = 2000bp.

To generate the first read, *R*_1_, in each model we first randomly selected the start position of the first nucleosome from the first 90bp of the read. The nucleosome occupancy status of each bp in the read was then simulated under the assumptions that each nucleosome occupies 147bp of DNA and that the length of linker DNA between nucleosomes can take any value between 10bp and 90bp with equal probability. For *i* = 1, …, 2000, we set *R*_1,*i*_ = 0 if base *i* is covered by a nucleosome and *R*_1,*i*_ = 1 otherwise.

The first model considers the case where nucleosomes are perfectly phased between molecules, i.e. nucleosomes are positioned in exactly the same place in each molecule. Each of the reads *R*_2_, …, *R*_200_ therefore has exactly the same nucleosome occupancy pattern as *R*_1_. In the second model, it is assumed that there is no phasing of nucleosomes between molecules, i.e. nucleosome positioning in one molecule is entirely independent of nucleosome positioning in other molecules. In this case, each read *R*_2_, …, *R*_200_ was simulated independently, using the same approach as used to simulate *R*_1_.

For each model, we used the simulated data to calculate single-molecule distance-dependent correlations in nucleosome-occupancy status. In particular, for *d* = 1, …, 500, the correlation in nucleosome occupancy status associated with bases a distance of *d*bps apart was calculated for read *i* via

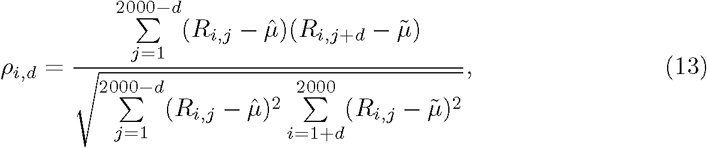

where

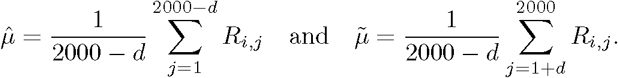

We then used the simulated data to derive bulk-level data associated with nucleosome occupancy. In particular, we calculated the bulk probability that base *j* ∈ {1, 2, …, 2000} is not occupied by a nucleosome or is occupied by a nucleosome, respectively, via

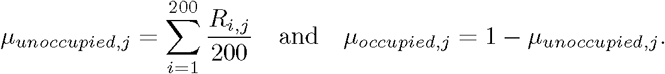

We then used this information to create “simulated” synthetic reads 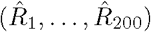 for each model in an analogous way as was done for the real data. Finally, we used the simulated synthetic reads to calculate distance-dependent correlations in nucleosome occupancy in a similar way as described above.

## Acknowledgments

We thank Chris Ponting, Marcus Wilson, Andreas Kapourani and members of the Sproul and Grima lab for helpful discussions about the manuscript.

LK is a cross-disciplinary post-doctoral fellow supported by funding from the University of Edinburgh and Medical Research Council (MC UU 00009/2). IK is funded by a CRUK PhD studentship and The A. G. Leventis Foundation Educational Grant. RG is supported by Leverhulme Trust research awards (RPG-2018-423 and RPG-2020-327). DS is a Cancer Research UK Career Development fellow (reference C47648/A20837), and work in his laboratory is also supported by an MRC university grant to the MRC Human Genetics Unit.

## Supporting information

**Fig. S1.**
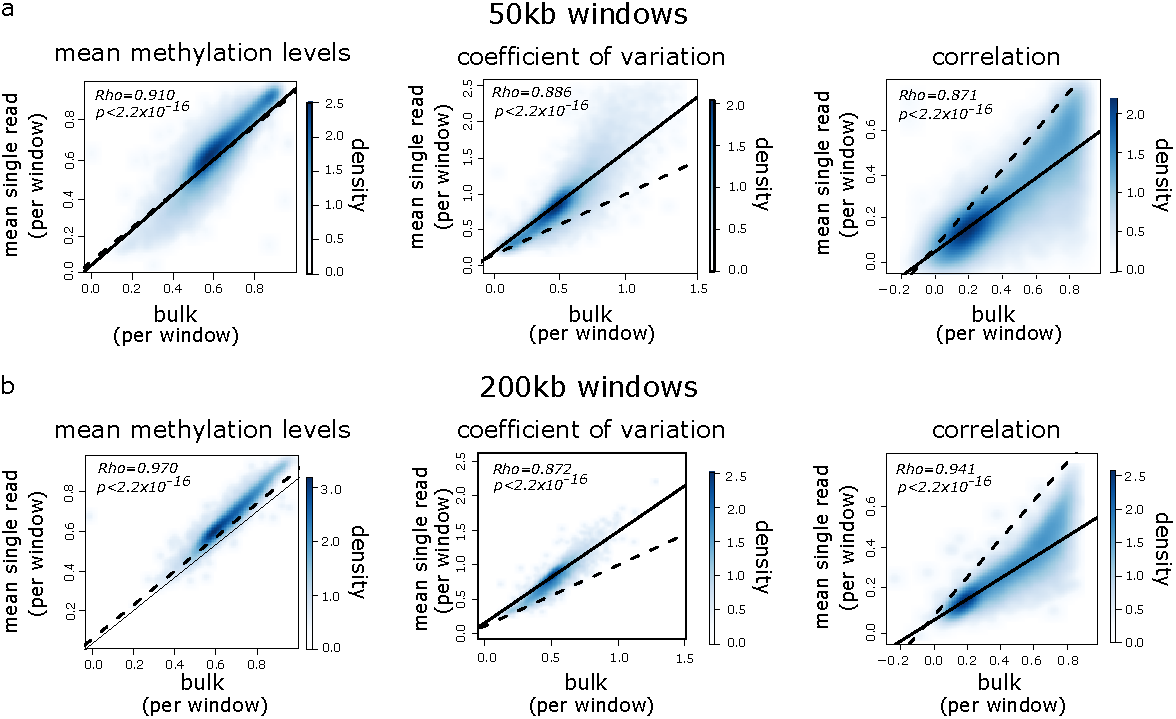
Large-scale methylation heterogeneity is underestimated by bulk analysis of methylomes independently of the window size used. Density scatter plots of mean single-read statistics vs. bulk statistics for 50kb genomic windows (*n* = 52, 840) (**a**) and 200kb genomic windows (*n* = 13, 439) (**b**). Left: mean methylation level, middle: coefficient of variation, right: correlation between neighbouring CpG sites. Dashed and solid lines show lines of identity and linear models fitted to the data respectively. Spearman correlations are shown, as are p-values from paired t-tests comparing bulk to single-read statistics.

**Fig. S2.**
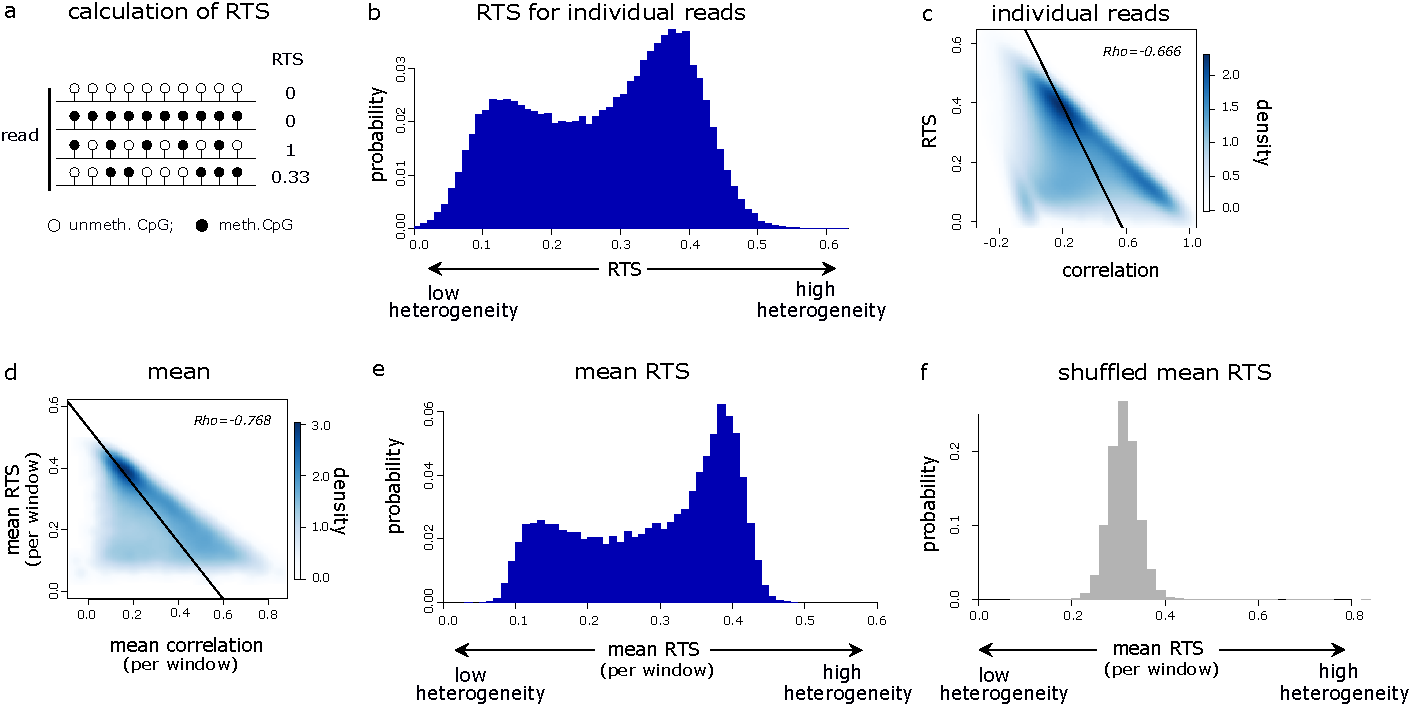
Intra-molecular heterogeneity varies across the human genome. **a** Schematic of RTS values calculated for hypothetical 10 CpG reads. **b** Histogram showing the RTS distribution for the 2, 908, 181 individual reads containing methylation information for ≥ 100 CpGs. **c** Density scatter plot of the RTS vs. correlation for individual reads. **d** Density scatter plot of the mean RTS vs. mean correlation for 100kb genomic windows. In **c, d** the solid line shows a linear model fitted to the data and the Spearman correlation coefficient is shown in the top right. **e** Histogram showing the distribution of mean RTS for 100kb windows (*n* = 26, 951) calculated using reads that align entirely within these windows. **f** Histogram showing the mean shuffled RTS in 100kb windows (*n* = 26, 951), calculated using random reads rather than those that align to each window.

**Fig. S3.**
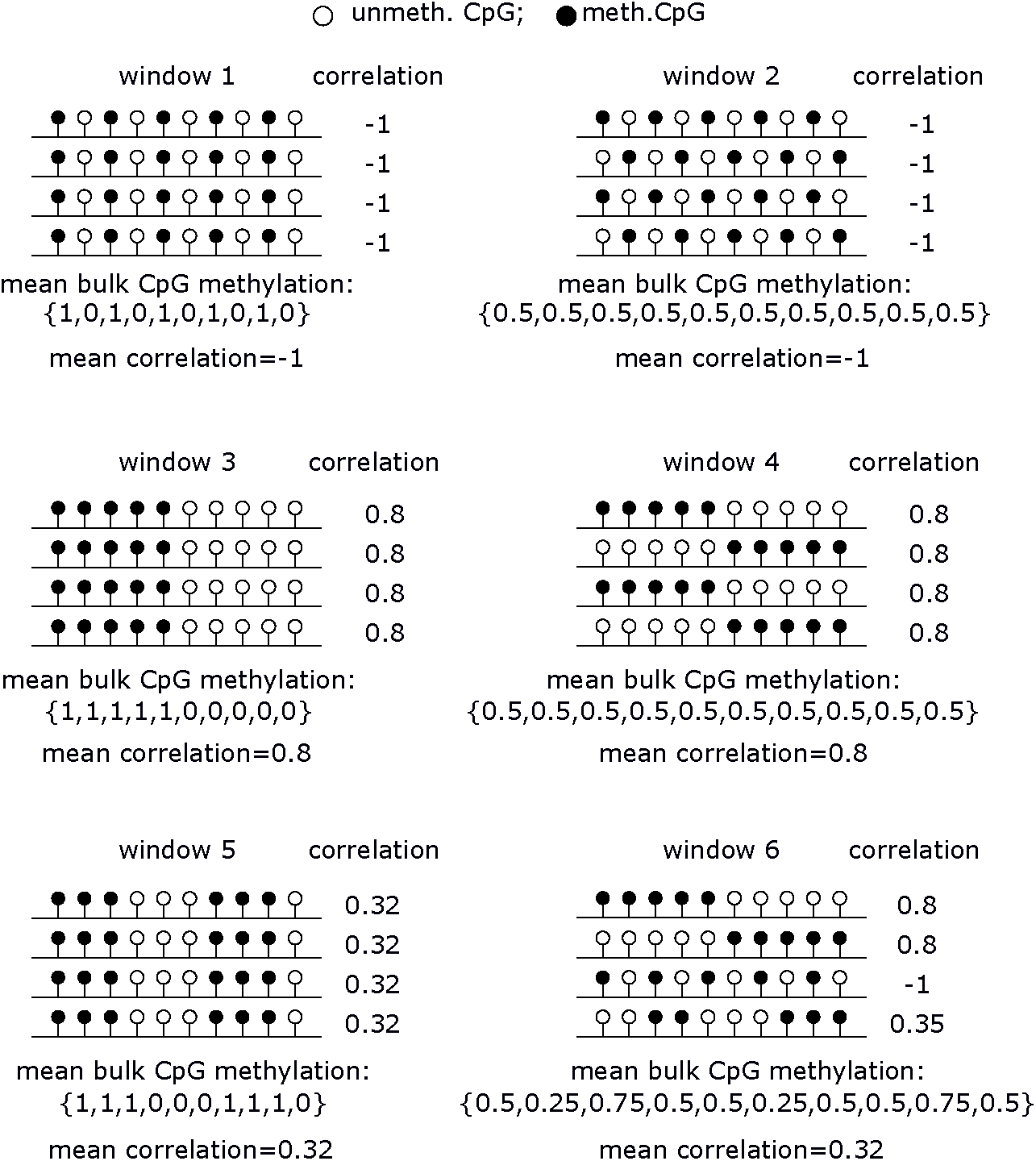
Equivalent levels of intra-molecular heterogeneity can be associated with different levels of inter-molecular heterogeneity. Shown are six hypothetical examples of genomic windows composed of 10 CpG sites and covered by four reads. For each example, the correlation associated with each read is shown to the right of the read and the mean correlation for the window is shown below each example. The mean bulk methylation of each individual CpG is indicated below the reads. Each pair of examples: windows 1 and 2, windows 3 and 4, windows 5 and 6, show situations where the same mean window correlation is associated with differing degrees of inter-molecular heterogeneity.

**Fig. S4.**
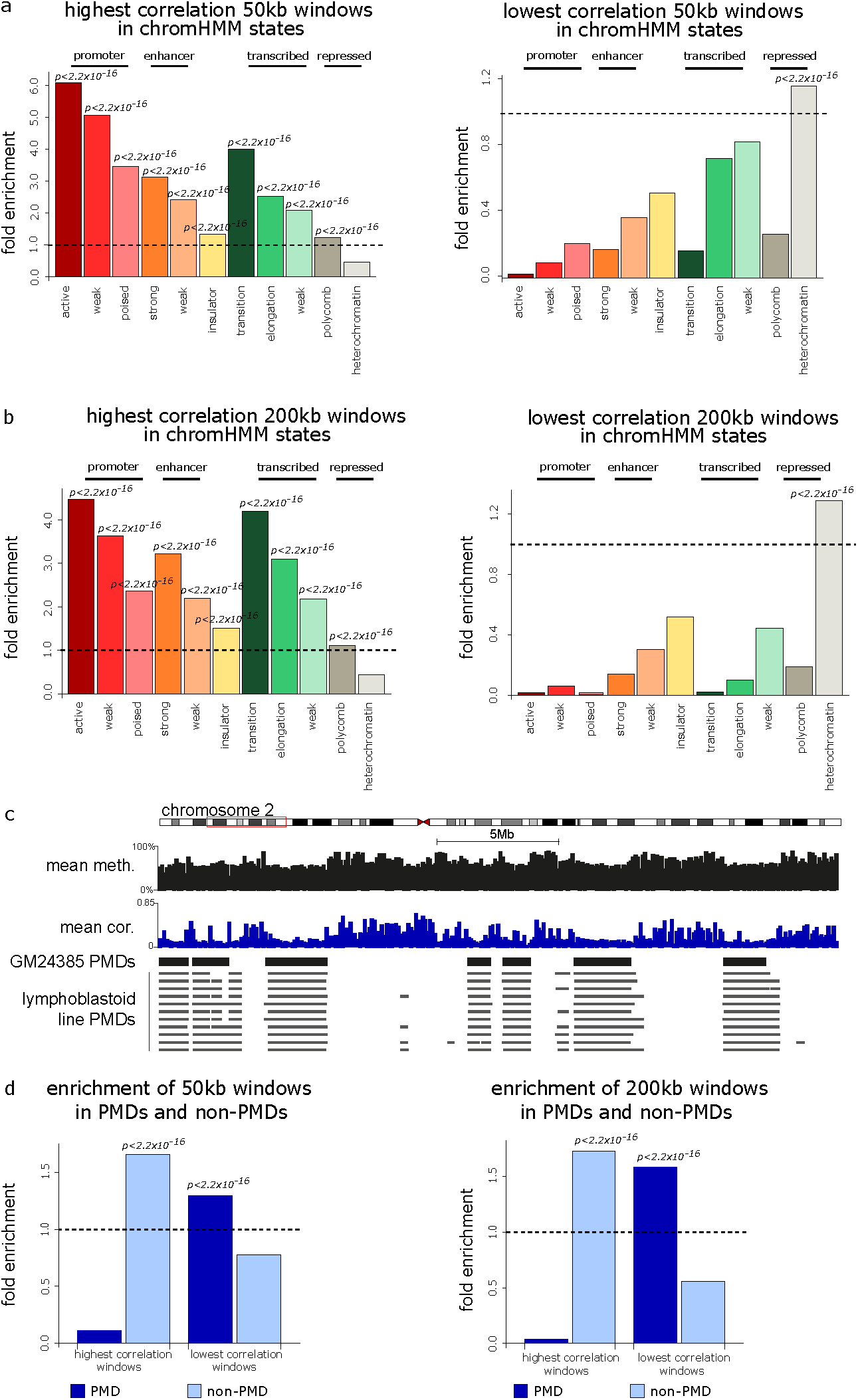
Heterogeneous genomic windows are enriched in heterochromatin and PMDs independently of the window size used. **a** Barplots showing enrichments of the least and most heterogeneous 50kb windows in GM12878 chromHMM states. **b** Barplots showing enrichment of the least and most heterogeneous 200kb windows in GM12878 chromHMM states. **c** Genome browser plot showing a representative genomic region with the mean DNA methylation and mean correlation in 100kb windows alongside GM24385 PMD locations and those previously identified in 11 other lymphoblastoid cell lines [25]. **d** Barplots showing enrichment of the least and most heterogeneous 50kb windows (left) and 200kb windows (right) in PMDs and non-PMDs. Shown are significant p-values from Wilcoxon tests.

**Fig. S5.**
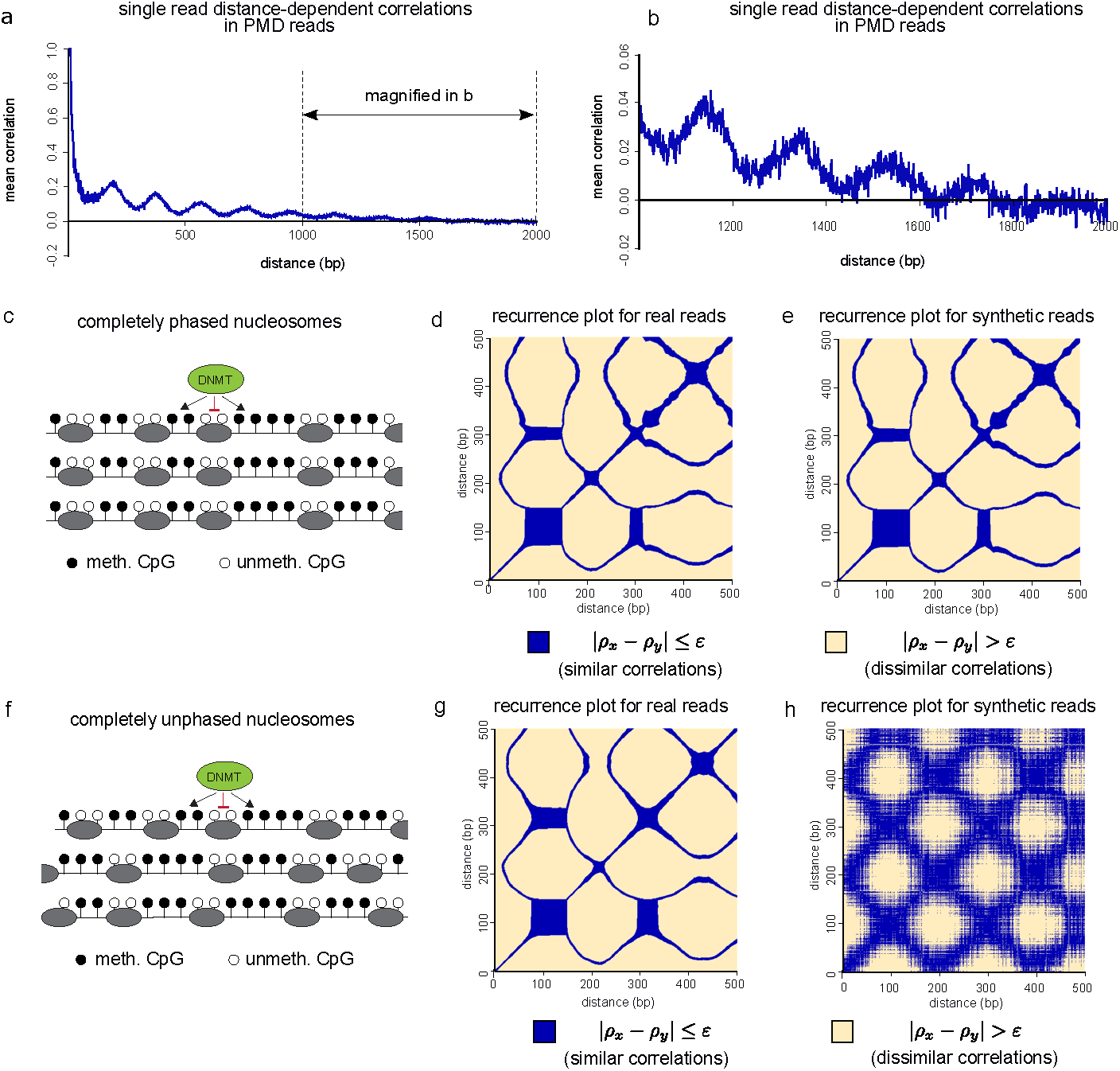
Periodic DNA methylation patterns are observed in PMDs within single molecules. **a** Plot of mean single-read distance-dependent correlation between CpG sites within 2kb of each other for 1, 007, 580 reads aligning entirely within PMDs. **B** Magnification of the region indicated in **a** (i.e. mean single-read distance-dependent correlation for distances 1kb-2kb in PMD reads). **c** Schematic of model of the interaction between DNA methylation and nucleosomes in the case of perfect nucleosomal phasing between molecules (see Methods). **d**,**e** single-molecule (**d**) and bulk (**e**) recurrence plots derived from the perfect phasing model. Here *ε* = 0.06 for comparison to recurrence plots associated with real data. **f** Model of completely unphased nucleosomes (Methods). Here it is assumed that nucleosomes are positioned randomly in each molecule. **g**,**h** Single-molecule (**g**) and bulk (**h**) recurrence plots derived from the completely unphased model. Again *ε* = 0.06 for comparison to recurrence plots associated with real data.

## References

1. Schübeler D. Function and information content of DNA methylation. Nature. 2015;517(7534):321–326.

2. Suzuki MM, Bird A. DNA methylation landscapes: provocative insights from epigenomics. Nature reviews genetics. 2008;9(6):465–476.

3. Lister R, Pelizzola M, Dowen RH, Hawkins RD, Hon G, Tonti-Filippini J, et al. Human DNA methylomes at base resolution show widespread epigenomic differences. nature. 2009;462(7271):315–322.

4. Urich MA, Nery JR, Lister R, Schmitz RJ, Ecker JR. MethylC-seq library preparation for base-resolution whole-genome bisulfite sequencing. Nature protocols. 2015;10(3):475–483.

5. Ziller MJ, Gu H, Müller F, Donaghey J, Tsai LTY, Kohlbacher O, et al. Charting a dynamic DNA methylation landscape of the human genome. Nature. 2013;500(7463):477–481.

6. Schultz MD, He Y, Whitaker JW, Hariharan M, Mukamel EA, Leung D, et al. Human body epigenome maps reveal noncanonical DNA methylation variation. Nature. 2015;523(7559):212–216.

7. Berman BP, Weisenberger DJ, Aman JF, Hinoue T, Ramjan Z, Liu Y, et al. Regions of focal DNA hypermethylation and long-range hypomethylation in colorectal cancer coincide with nuclear lamina–associated domains. Nature genetics. 2012;44(1):40.

8. Landan G, Cohen NM, Mukamel Z, Bar A, Molchadsky A, Brosh R, et al. Epigenetic polymorphism and the stochastic formation of differentially methylated regions in normal and cancerous tissues. Nature genetics. 2012;44(11):1207–1214.

9. Sheffield NC, Pierron G, Klughammer J, Datlinger P, Schönegger A, Schuster M, et al. DNA methylation heterogeneity defines a disease spectrum in Ewing sarcoma. Nature medicine. 2017;23(3):386–395.

10. Jenkinson G, Pujadas E, Goutsias J, Feinberg AP. Potential energy landscapes identify the information-theoretic nature of the epigenome. Nature genetics. 2017;49(5):719–729.

11. Shipony Z, Mukamel Z, Cohen NM, Landan G, Chomsky E, Zeliger SR, et al. Dynamic and static maintenance of epigenetic memory in pluripotent and somatic cells. Nature. 2014;513(7516):115–119.

12. Li S, Garrett-Bakelman FE, Chung SS, Sanders MA, Hricik T, Rapaport F, et al. Distinct evolution and dynamics of epigenetic and genetic heterogeneity in acute myeloid leukemia. Nature medicine. 2016;22(7):792–799.

13. Abante J, Fang Y, Feinberg A, Goutsias J. Detection of haplotype-dependent allele-specific DNA methylation in WGBS data. Nature communications. 2020;11(1):1–13.

14. Bergstedt J, Azzou SAK, Tsuo K, Jaquaniello A, Urrutia A, Rotival M, et al. The immune factors driving DNA methylation variation in human blood. Nature communications. 2022;13(1):1–20.

15. Luo C, Keown CL, Kurihara L, Zhou J, He Y, Li J, et al. Single-cell methylomes identify neuronal subtypes and regulatory elements in mammalian cortex. Science. 2017;357(6351):600–604.

16. Tanaka K, Okamoto A. Degradation of DNA by bisulfite treatment. Bioorganic & medicinal chemistry letters. 2007;17(7):1912–1915.

17. Payne A, Holmes N, Rakyan V, Loose M. BulkVis: a graphical viewer for Oxford nanopore bulk FAST5 files. Bioinformatics. 2019;35(13):2193–2198.

18. Simpson JT, Workman RE, Zuzarte P, David M, Dursi L, Timp W. Detecting DNA cytosine methylation using nanopore sequencing. Nature methods. 2017;14(4):407–410.

19. Scherer M, Nebel A, Franke A, Walter J, Lengauer T, Bock C, et al. Quantitative comparison of within-sample heterogeneity scores for DNA methylation data. Nucleic acids research. 2020;48(8):e46–e46.

20. Robinson JT, Thorvaldsdóttir H, Winckler W, Guttman M, Lander ES, Getz G, et al. Integrative genomics viewer. Nature biotechnology. 2011;29(1):24–26.

21. Hetzel S, Giesselmann P, Reinert K, Meissner A, Kretzmer H. RLM: fast and simplified extraction of read-level methylation metrics from bisulfite sequencing data. Bioinformatics. 2021;37(21):3934–3935.

22. Consortium EP, et al. An integrated encyclopedia of DNA elements in the human genome. Nature. 2012;489(7414):57.

23. Ernst J, Kheradpour P, Mikkelsen TS, Shoresh N, Ward LD, Epstein CB, et al. Mapping and analysis of chromatin state dynamics in nine human cell types. Nature. 2011;473(7345):43–49.

24. Hovestadt V, Jones DT, Picelli S, Wang W, Kool M, Northcott PA, et al. Decoding the regulatory landscape of medulloblastoma using DNA methylation sequencing. Nature. 2014;510(7506):537–541.

25. Decato BE, Qu J, Ji X, Wagenblast E, Knott SR, Hannon GJ, et al. Characterization of universal features of partially methylated domains across tissues and species. Epigenetics & chromatin. 2020;13(1):1–14.

26. Gaidatzis D, Burger L, Murr R, Lerch A, Dessus-Babus S, Schübeler D, et al. DNA sequence explains seemingly disordered methylation levels in partially methylated domains of Mammalian genomes. PLoS genetics. 2014;10(2):e1004143.

27. Zhang L, Xie WJ, Liu S, Meng L, Gu C, Gao YQ. DNA methylation landscape reflects the spatial organization of chromatin in different cells. Biophysical journal. 2017;113(7):1395–1404.

28. Berman BP, Liu Y, Kelly TK. DNA methylation marks inter-nucleosome linker regions throughout the human genome. PeerJ PrePrints; 2013.

29. Marwan N, Romano MC, Thiel M, Kurths J. Recurrence plots for the analysis of complex systems. Physics reports. 2007;438(5-6):237–329.

30. Giuliani A, Manetti C. Hidden peculiarities in the potential energy time series of a tripeptide highlighted by a recurrence plot analysis: a molecular dynamics simulation. Physical Review E. 1996;53(6):6336.

31. Manetti C, Giuliani A, Ceruso MA, Webber Jr CL, Zbilut JP. Recurrence analysis of hydration effects on nonlinear protein dynamics: multiplicative scaling and additive processes. Physics Letters A. 2001;281(5-6):317–323.

32. Van Holde KE. Chromatin. Springer Science & Business Media; 2012.

33. Baldi S, Korber P, Becker PB. Beads on a string—nucleosome array arrangements and folding of the chromatin fiber. Nature structural & molecular biology. 2020;27(2):109–118.

34. Teif VB, Vainshtein Y, Caudron-Herger M, Mallm JP, Marth C, Höfer T, et al. Genome-wide nucleosome positioning during embryonic stem cell development. Nature structural & molecular biology. 2012;19(11):1185–1192.

35. Valouev A, Johnson SM, Boyd SD, Smith CL, Fire AZ, Sidow A. Determinants of nucleosome organization in primary human cells. Nature. 2011;474(7352):516–520.

36. Felle M, Hoffmeister H, Rothammer J, Fuchs A, Exler JH, Längst G. Nucleosomes protect DNA from DNA methylation in vivo and in vitro. Nucleic acids research. 2011;39(16):6956–6969.

37. Schrader A, Gross T, Thalhammer V, Längst G. Characterization of Dnmt1 binding and DNA methylation on nucleosomes and nucleosomal arrays. PloS one. 2015;10(10):e0140076.

38. Robertson AK, Geiman TM, Sankpal UT, Hager GL, Robertson KD. Effects of chromatin structure on the enzymatic and DNA binding functions of DNA methyltransferases DNMT1 and Dnmt3a in vitro. Biochemical and biophysical research communications. 2004;322(1):110–118.

39. Takeshima H, Suetake I, Shimahara H, Ura K, Tate Si, Tajima S. Distinct DNA methylation activity of Dnmt3a and Dnmt3b towards naked and nucleosomal DNA. Journal of biochemistry. 2006;139(3):503–515.

40. Takeshima H, Suetake I, Tajima S. Mouse Dnmt3a preferentially methylates linker DNA and is inhibited by histone H1. Journal of molecular biology. 2008;383(4):810–821.

41. Kelly TK, Liu Y, Lay FD, Liang G, Berman BP, Jones PA. Genome-wide mapping of nucleosome positioning and DNA methylation within individual DNA molecules. Genome research. 2012;22(12):2497–2506.

42. Lee I, Razaghi R, Gilpatrick T, Molnar M, Gershman A, Sadowski N, et al. Simultaneous profiling of chromatin accessibility and methylation on human cell lines with nanopore sequencing. Nature Methods. 2020;17(12):1191–1199.

43. Gaffney DJ, McVicker G, Pai AA, Fondufe-Mittendorf YN, Lewellen N, Michelini K, et al. Controls of nucleosome positioning in the human genome. PLoS genetics. 2012;8(11):e1003036.

44. Baldi S, Krebs S, Blum H, Becker PB. Genome-wide measurement of local nucleosome array regularity and spacing by nanopore sequencing. Nature structural & molecular biology. 2018;25(9):894–901.

45. Lai B, Gao W, Cui K, Xie W, Tang Q, Jin W, et al. Principles of nucleosome organization revealed by single-cell micrococcal nuclease sequencing. Nature. 2018;562(7726):281–285.

46. Abdulhay NJ, McNally CP, Hsieh LJ, Kasinathan S, Keith A, Estes LS, et al. Massively multiplex single-molecule oligonucleosome footprinting. Elife. 2020;9:e59404.

47. Orphanides G, LeRoy G, Chang CH, Luse DS, Reinberg D. FACT, a factor that facilitates transcript elongation through nucleosomes. Cell. 1998;92(1):105–116.

48. Zhou W, Dinh HQ, Ramjan Z, Weisenberger DJ, Nicolet CM, Shen H, et al. DNA methylation loss in late-replicating domains is linked to mitotic cell division. Nature genetics. 2018;50(4):591–602.

49. Endicott JL, Nolte PA, Shen H, Laird PW. Cell division drives DNA methylation loss in late-replicating domains in primary human cells. Nature Communications. 2022;13(1):6659.

50. Ming X, Zhang Z, Zou Z, Lv C, Dong Q, He Q, et al. Kinetics and mechanisms of mitotic inheritance of DNA methylation and their roles in aging-associated methylome deterioration. Cell research. 2020;30(11):980–996.

51. Bar D, Fishman L, Zheng Y, Unterman I, Schlesinger D, Eden A, et al. A local sequence signature defines a subset of heterochromatin-associated CpGs with minimal loss of methylation in healthy tissues but extensive loss in cancer. bioRxiv. 2022; p. 2022–08.

52. Struhl K, Segal E. Determinants of nucleosome positioning. Nature structural & molecular biology. 2013;20(3):267–273.

53. Jeltsch A, Adam S, Dukatz M, Emperle M, Bashtrykov P. Deep enzymology studies on DNA methyltransferases reveal novel connections between flanking sequences and enzyme activity. Journal of molecular biology. 2021;433(19):167186.

54. Landau DA, Clement K, Ziller MJ, Boyle P, Fan J, Gu H, et al. Locally disordered methylation forms the basis of intratumor methylome variation in chronic lymphocytic leukemia. Cancer cell. 2014;26(6):813–825.

55. Guo S, Diep D, Plongthongkum N, Fung HL, Zhang K, Zhang K. Identification of methylation haplotype blocks aids in deconvolution of heterogeneous tissue samples and tumor tissue-of-origin mapping from plasma DNA. Nature genetics. 2017;49(4):635–642.

56. Quinlan AR, Hall IM. BEDTools: a flexible suite of utilities for comparing genomic features. Bioinformatics. 2010;26(6):841–842.

57. Technologies ON. Guppy protocol: modified base calling;.

58. Li H. Minimap2: pairwise alignment for nucleotide sequences. Bioinformatics. 2018;34(18):3094–3100.

59. Liu Y, Rosikiewicz W, Pan Z, Jillette N, Wang P, Taghbalout A, et al. DNA methylation-calling tools for Oxford Nanopore sequencing: a survey and human epigenome-wide evaluation. Genome biology. 2021;22(1):1–33.

60. Akbari V, Garant JM, O’Neill K, Pandoh P, Moore R, Marra MA, et al. Megabase-scale methylation phasing using nanopore long reads and NanoMethPhase. Genome biology. 2021;22(1):1–21.

61. Neph S, Kuehn MS, Reynolds AP, Haugen E, Thurman RE, Johnson AK, et al. BEDOPS: high-performance genomic feature operations. Bioinformatics. 2012;28(14):1919–1920.

62. Frankish A, Diekhans M, Jungreis I, Lagarde J, Loveland JE, Mudge JM, et al. GENCODE 2021. Nucleic acids research. 2021;49(D1):D916–D923.

63. Illingworth RS, Gruenewald-Schneider U, Webb S, Kerr AR, James KD, Turner DJ, et al. Orphan CpG islands identify numerous conserved promoters in the mammalian genome. PLoS genetics. 2010;6(9):e1001134.

64. Masalmeh RHA, Taglini F, Rubio-Ramon C, Musialik KI, Higham J, Davidson-Smith H, et al. De novo DNA methyltransferase activity in colorectal cancer is directed towards H3K36me3 marked CpG islands. Nature communications. 2021;12(1):1–13.

